# A Causal Model of Ion Interference Enables Assessment and Correction of Ratio Compression in Multiplex Proteomics

**DOI:** 10.1101/2022.06.24.497446

**Authors:** Moritz Madern, Wolfgang Reiter, Florian Stanek, Natascha Hartl, Karl Mechtler, Markus Hartl

## Abstract

Multiplex proteomics using isobaric labeling tags has emerged as a powerful tool for the simultaneous relative quantification of peptides and proteins across multiple experimental conditions. However, the quantitative accuracy of the approach is largely compromised by ion interference, a phenomenon that causes fold changes to appear compressed. The degree of compression is generally unknown, and the contributing factors are poorly understood. In this study, we thoroughly characterized ion interference at the MS2 level using a defined two-proteome experimental system with known ground-truth. We discovered remarkably poor agreement between the apparent precursor purity in the isolation window and the actual level of observed reporter ion interference in MS2-scans – a discrepancy that we found resolved by considering co-fragmentation of peptide ions hidden within the spectral “noise” of the MS1 isolation window. To address this issue, we developed a regression modeling strategy to accurately predict estimates of reporter ion interference in any dataset. Finally, we demonstrate the utility of our procedure for improved fold change estimation and unbiased PTM site-to-protein normalization. All computational tools and code required to apply this method to any MS2 TMT dataset are documented and freely available.

## INTRODUCTION

In mass spectrometry (MS)-based proteomics, isobaric labeling systems such as Tandem Mass Tags (TMT) have been established to efficiently quantify proteins across multiple conditions within a single experiment^1–3^. Their high throughput is based on the ability to combine up to 18 samples in one experiment^4^, using sample-specific isobaric labels that allow joint processing and measurement of peptides. Upon fragmentation of the labels in the mass spectrometer, sample-specific reporter ions are generated^1^, revealing the relative abundance of peptides and proteins for each sample. As all samples are measured at the same time, the approach is largely free of missing values. Further, more complex sample processing steps like offline-fractionation^2^ or the enrichment for post-translational modifications (PTMs)^2, 3^ benefit from the early pooling of samples by eliminating additional technical variation.

However, despite the elegance in design, multiplex proteomics struggles with accurate relative quantification due to the phenomenon termed ion interference. Concomitant fragmentation of targeted precursor ions along with interfering non-precursor ions in the same isolation window results in convoluted reporter ion profiles^5, 6^. As the majority of proteins in a typical large-scale proteomic experiment are non-differentially expressed, relative quantitative differences of precursor peptides become suppressed by a uniform background of interfering reporter ion signal. This effect is also referred to as “ratio compression”^6^. Extensive sample fractionation and decreased isolation window widths help to reduce this effect^7–10^, yet they do not completely remove it. Interestingly, despite ratio compression, the ability to distinguish differentially from non-differentially expressed proteins using isobaric labeling-based quantification is comparable or even outperforms label free quantification (LFQ) due to increased quantitative precision and data completeness within the same multiplex set^11^. However, compressed fold changes make it more difficult to interpret results in a biologically meaningful way. Therefore, different strategies have been developed to address and mitigate the ratio compression problem.

Promising technical solutions encompass gas-phase purification^12^, dual isolation width acquisition^9^, High-Field Asymmetric-Waveform Ion Mobility MS (FAIMS)^13^ and MS3-based quantification^10, 14–16^. Unfortunately, these methods come with their own set of limitations as they require specific instrumentation for data acquisition^10, 12–15^, and the measurement of additional scans per precursor generally decreases measurement speed and thus the number of IDs^9, 10, 14–16^. New isobaric tags that generate interference-free reporter ions have been developed by Virreira Winter et al.^17^ but they are commercially not yet available, preventing wider testing and adoption by other labs. Wühr *et al*. described an alternative quantification method in which the complementary fragment ion cluster in the MS2 spectrum is used for the quantitative profiling^18, 19^. Unfortunately, most current MS instruments lack the required resolution to separate heavy N and C isotope mass differences in the higher m/z region of the complementary ions, which reduces the multiplexing capabilities to a maximum of 9 channels^4, 20^. Another hurdle of quantification via the complementary ion cluster lies in the complementary ion formation itself, as currently available isobaric labeling systems are not optimized to generate complementary fragment ions^21^. A recent study examined the use of combined 11plex and 18plex TMT reagents to estimate interference levels^22^. These estimates successfully enabled the decompression of fold changes; however, the general applicability of this innovative multiplexing strategy needs to be further evaluated.

Other strategies address ratio compression *in silico*. Savitski *et al*. proposed an interference correction algorithm based on the precursor ion purity in the MS1 isolation window range^23^. Niu *et al*. estimated the level of interference via Lys-y1 and Arg-y1 ion intensities in MS2-scans, ascribing them to either target or interfering peptides^8^. Other approaches attempt to correct ratio compression by modeling the quantitative relationship between observed and theoretical fold changes provided by a synthetic peptide spike-in^6, 24^. Similarly, modeling observed fold changes in dependence of precursor ion purities and signal to noise ratios for different peptides of the same protein has been proposed^25^. Altogether, we believe that these strategies have certain limitations due to their phenomenological approach in addressing the interference issue. They rely on correlations between ion interference and a single measured variable, providing only a partial explanation for the extent of ratio compression in an experiment. This simplified perspective can be attributed to the current limitations in our understanding of the causal factors underlying interference. Consequently, the proposed computational methods are not fully equipped to address ratio compression at the level of individual PSMs, thereby restricting their effectiveness in comprehensively mitigating ratio compression throughout an entire experiment.

In this study, we aimed to address ratio compression by building on a holistic mechanistic understanding of the underlying phenomenon itself, i.e. ion interference. To this end, we designed a controlled two-proteome experiment which allowed us to directly observe reporter ion interference (here defined as the reporter ion signal at MS2 level caused by ion interference) for a substantial fraction of peptides while simultaneously monitoring the accompanied ratio compression effect on known theoretical fold changes. Taking advantage of this design, we first explored the dependence of ion interference on varying measurement parameters and the accompanying effect on differential expression (DE) analysis. We then aimed to causally understand the measured reporter ion interference by means of multiple linear regression modeling. We found that ion interference is only duly explained when accounting for spectral “noise” at the MS1 level, as the apparent ion purity in the isolation window generally underestimates the true extent of reporter ion interference at MS2 level. By generalizing such a model, we found a way to accurately predict reporter ion interference at the PSM-level in any MS2-quantified multiplex proteomics dataset. Finally, we demonstrate how this gain in information can be used to decompress fold changes affected by ratio compression, and carry out normalization of PTM site to underlying protein abundances unbiased by differences in ion interference.

## EXPERIMENTAL PROCEDURES

### Experimental design and statistical rationale

We conducted a two-proteome multiplex (TMTpro 16plex) proteomics experiment, following a similar approach to previous studies^8–11, 13–16, 24–29^. The experiment involved proteomes from human Jurkat E6-1 cells and the budding yeast strain W303-1A (**Figure 1**). Human peptides constituted the majority of each sample and served as a stable quantitative background. Yeast peptides of comparatively low abundance were varied in defined ratios across four groups of three technical replicates each, allowing for fold change calculation and differential expression testing. Three reporter ion channels (128N, 129N, and 130N) were completely free of yeast peptides, thereby providing a direct quantitative readout of human-derived reporter ion interference at the MS2 level. This setup allowed us to explore and predict reporter ion interference at the MS2 level using multiple linear regression modeling, as further explained in the subsequent sections.

**Figure 1.**
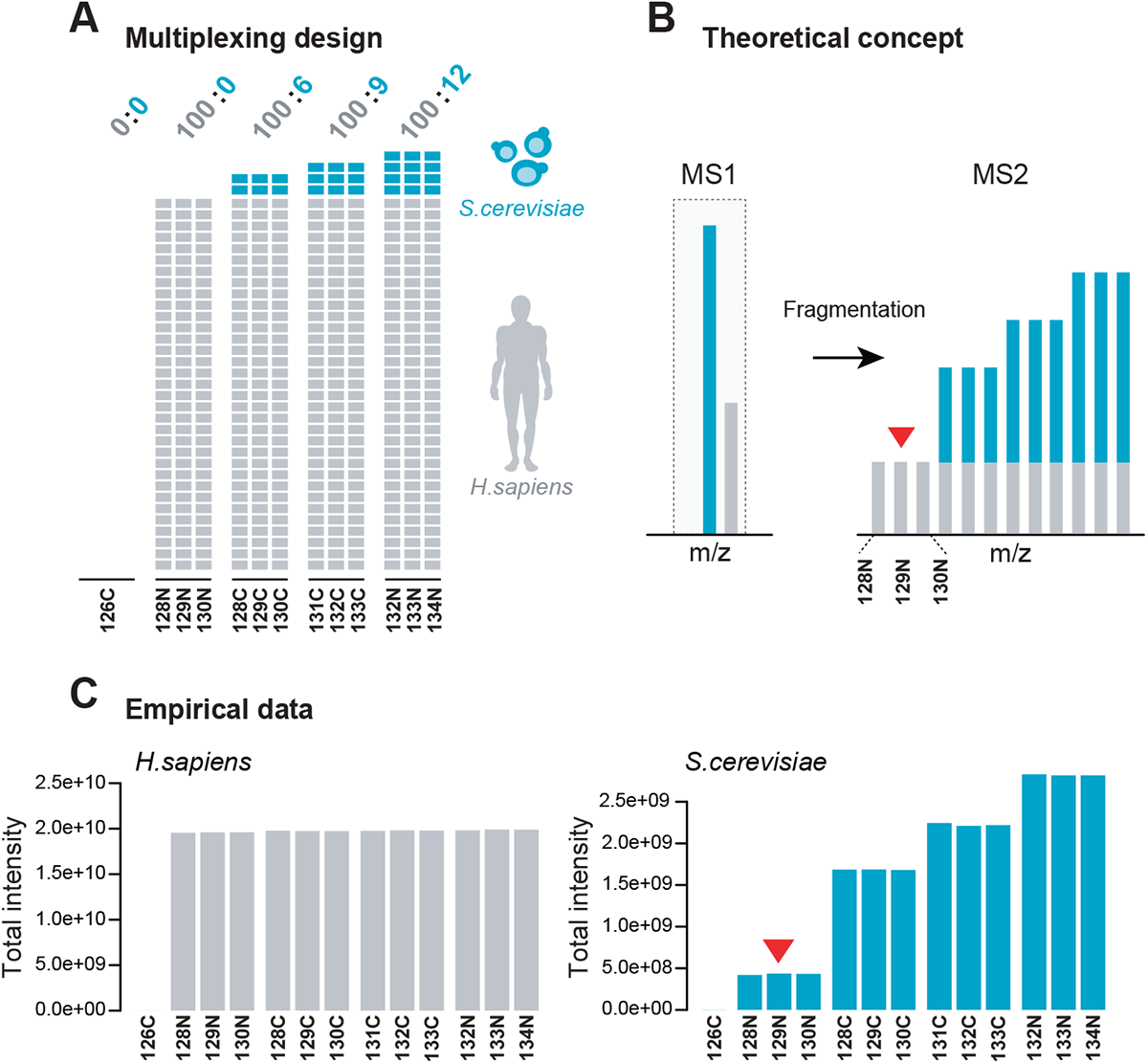
Artificial two-proteome dataset for exploring ion interference. **(A)** Schematic depiction of sample preparation and TMTpro 16plex labeling scheme. Samples were defined mixtures of *H. sapiens* Jurkat (grey) and *S. cerevisiae* (turquoise) peptides from whole-proteome digests. Relative mass abundance ratios (w/w) of human to yeast are displayed on top of each sample group. **(B)** Concept of measuring reporter ion interference at the MS2 level via channels 128N, 129N and 130N (marked by red arrow) for a targeted yeast peptide. (**C)** Total measured channel intensities of quantified human (left) and yeast (right) peptides after between-sample normalization. The red arrow marks human-derived reporter ion interference in yeast peptide measurements.

### Cell culture

*S. cerevisiae* yeast cells W303-1A were cultured shaking (200 rpm) at 30°C until mid-log phase (OD600nm ∼1) in 50 ml rich medium (YPD; 1 % yeast extract, 2 % peptone, 2 % glucose). Cells were harvested by filtration (Protran 0.45 μm nitrocellulose membrane, Amersham) and frozen in liquid nitrogen. Human Jurkat E6-1 cells were cultured in Roswell Park Memorial Institute (RPMI) 1640 media (Fibco, Grand Island, NY), supplemented with 10 % fetal bovine serum (Gibco), penicillin and streptomycin (Gibco). Cells were pelleted and washed with 1 x phosphate buffered saline (PBS) prior to freezing in liquid nitrogen.

### Proteomic sample preparation

Frozen yeast cell pellets were resuspended in lysis buffer (8 M urea, 50 mM Tris buffer pH 8.0, 150 mM NaCl, 1 mM PMSF, 5 mM sodium butyrate, benzonase and protease inhibitor cocktail) and subsequently lysed by bead-beating (FastPrep®-24 device, MP Biomedicals). Frozen Jurkat cells were dissolved in lysis buffer and lysed via sonication. After removal of insoluble debris by centrifugation at 16,000 x g at 4°C for 10 min, yeast and human proteins were precipitated by the addition of three times the volume of cold acetone and subsequently incubated overnight at -20°C. Proteins were pelleted by centrifugation at 15,000 x g, dissolved in 8 M Urea, 50 mM ammonium bicarbonate (ABC) buffer and treated with 10 mM DTT (Dithiothreitol) for 45 min to reduce protein disulfide bonds. Alkylation of reduced cysteines was performed by adding iodoacetamide (IAA) to a final concentration of 20 mM, followed by incubation for 30 min in the dark. The remaining IAA was quenched and samples were diluted to 4 M urea for digestion with Lys-C 1:50 (w/w). After incubation for 2 h at room temperature, samples were further diluted to a final concentration of 1 M urea, and digestion with trypsin 1:50 (w/w) was carried out overnight at 37°C. Finally, tryptic peptides were desalted using SepPak C18 cartridges (Waters), and eluates were shortly vacuum centrifuged and subsequently lyophilized.

### Two-proteome controlled experiment

Lyophilized tryptic yeast and human peptides were reconstituted in 0.1% trifluoroacetic acid (TFA) to reach a peptide concentration of 3 μg/μl each. Four distinct artificial two-proteome mixtures were created as follows: A constant amount of human peptide solution (300 μl) was initially prepared for each mixture. Then, a comparatively small, variable amount of yeast peptide solution was added to each solution, namely 0, 18, 27 and 36 μl, respectively – resulting in theoretical fold changes of 1.33, 1.5 and 2.0. The yeast-human mixtures were subsequently lyophilized and again reconstituted in 120 μl 100 mM TEAB (Sigma) each. Next, each mixture was split into 3 technical replicates, resulting in 12 peptide samples in total. TMT labeling was conducted according to a labeling scheme that minimizes isotopic interference across groups (**Figure 1A**). Additionally, a thirteenth sample without peptides was labeled in parallel using TMT label 126C to interrogate the effect of hydrolyzed and quenched label on ion interference.

### TMT-labeling

Dried TMTpro 16plex 0.5 mg label reagents (Thermo Fisher) were resuspended in 30 μl anhydrous acetonitrile (ACN) and mixed with 30 μl 100 mM TEAB peptide solution, resulting in a ratio of TMT label to peptide of approximately 2:1 (w/w); and a final peptide concentration of about 4.2 μg/μl per 60 μl labeling reaction. After incubation for 1 h, a small aliquot of each reaction was combined in 100 μl of 0.1 % TFA. This pooled sample was measured on the mass spectrometer to ascertain a labeling efficiency of > 99 % in each channel by using an in-house R script for the analysis. The labeling reactions were subsequently quenched by addition of 7 μl of 5 % hydroxylamine and incubation for 25 min at 400 rpm at RT. Finally, all 13 labeled samples were combined 1:1, desalted via SepPak C18 cartridges, vacuum centrifuged and lyophilized prior to neutral pH fractionation.

### Neutral pH fractionation

Dried labeled peptides were reconstituted in 100 μl 10 mM ammonium formate pH 6.8. After sonication, the solution was injected into UltiMate 3000 Dual LC pHPLC System equipped with a C18 column (xBridge Peptide BEH C18, 25 cm x 4.6 mm, 3.5 µm, Waters). 1 ml fractions were collected during separation on a 5-50% ACN 1 ml/min gradient in 10 mM ammonium formate buffer pH 6.8 at a flow rate of 1 ml/min. The resulting 1 ml fractions were vacuum centrifuged to evaporate the ACN, and then pooled according to a cyclical pooling scheme^30^ to create six sample pools of reduced complexity (labelled P1-P6). The pools were desalted via SepPak C18 cartridges and stored at -80°C until measurement on the LC-MS system.

### Acetylated peptide enrichment

Acetylomes were purified using the PTMScan Acetyl-Lysine Motif Kit (Cell Signaling). Roughly 600µg of peptides were dissolved in 175 µl of 1 x PTMScan IAP buffer. An appropriate amount of antibody bead slurry was washed four times with PBS, added to the peptide solution and incubated for two hours at 4 °C with end-over-end rotation. The beads were subsequently washed twice in 1 x IAP and 3 x with HPLC-grade water (Fisher Chemical). Peptides were eluted with 2 x 20 μl of 0.15% TFA. Eluates were united and desalted using C18 StageTips^31^

### Phosphorylated peptide enrichment using TiO2

An aliquot (peptide: TiO2 resin = 1 : 6) of TiO2 (Titansphere TiO, GL Sciences, 5020-75000) was washed twice with 50% methanol (Fisher chemical, A456-212) and twice with glycolic acid solution (1 M glycolic acid (Sigma Aldrich, 124737-25G), 70% ACN (VWR, 83639.320), 3% TFA (Thermo Scientific, 28903)). Peptides were dissolved in glycolic acid solution, mixed with the TiO2 resin, incubated rotating at room temperature for 30 min, transferred onto Mobicol columns (MoBiTec, M1003, M2110) and shortly centrifuged in a table centrifuge to remove unphosphorylated peptides. The resin was washed twice with glycolic acid solution, twice with 200 µl 70%, ACN 3%, TFA and twice with 1% ACN, 0,1% TFA. Phosphorylated peptides were eluted twice using 150 µl 300 mM ammonium hydroxide (VWR, 1.05432.1000), eluates were united and immediately acidified with conc. TFA to a pH of 2.5. Samples were desalted using a standard C18 StageTip protocol^31^.

### Measurement on the LC-MS system

Unless specified otherwise, the results are based on data generated from the yeast-human fraction pools of reduced sample complexity (labelled P1-P6) measured via MS2-based quantification. In brief, for each of the six pools, a total of 200 ng peptide material was measured on a Q Exactive HF-X Orbitrap mass spectrometer (Thermo Fisher), coupled to the LC-system with a nano-spray ion-source using coated emitter tips (PepSep, MSWil). Peptides were separated along a segmented 2 h 2-40% ACN gradient on the Ultimate 3000 RSLC nano-flow chromatography system (Thermo-Fisher) using a pre-column for sample loading (Acclaim PepMap C18, 2 cm × 0.1 mm, 5 μm, Thermo-Fisher), and a C18 analytical column (Acclaim PepMap C18, 50 cm × 0.75 mm, 2 μm, Thermo-Fisher) for separation, at a flow rate of 230 nl/min. On the mass spectrometer, the following instrument settings were applied. For MS1 scans: resolution 120 k, target AGC 1e6, maximum IT 60 ms; for MS2 scans: resolution 45 k, target AGC 2e5, maximum IT 120 ms, respectively. Precursor ions were targeted for fragmentation in a top 20 method using a dynamic exclusion time of 20 s. Target ions were isolated at an isolation window width of 0.7 Th and fragmented at NCE of 35. Precursor ion charge states of 1 or >7 were excluded.

Measurements for the interrogation of varying measurement-specific parameters (**Figure 2**) were performed on Q-Exactive HF-X (Thermo Fisher) mass spectrometers with generally similar instrument settings – with one exception: For the comparison of varying quantifications strategies, all measurements were performed on an Orbitrap Eclipse Tribid mass spectrometer (Thermo Fisher), as described in detail in **Table S1**. To the best of our abilities, the involved instrument settings were set as to best optimize the performance of each individual quantification method while ensuring maximum comparability.

**Figure 2.**
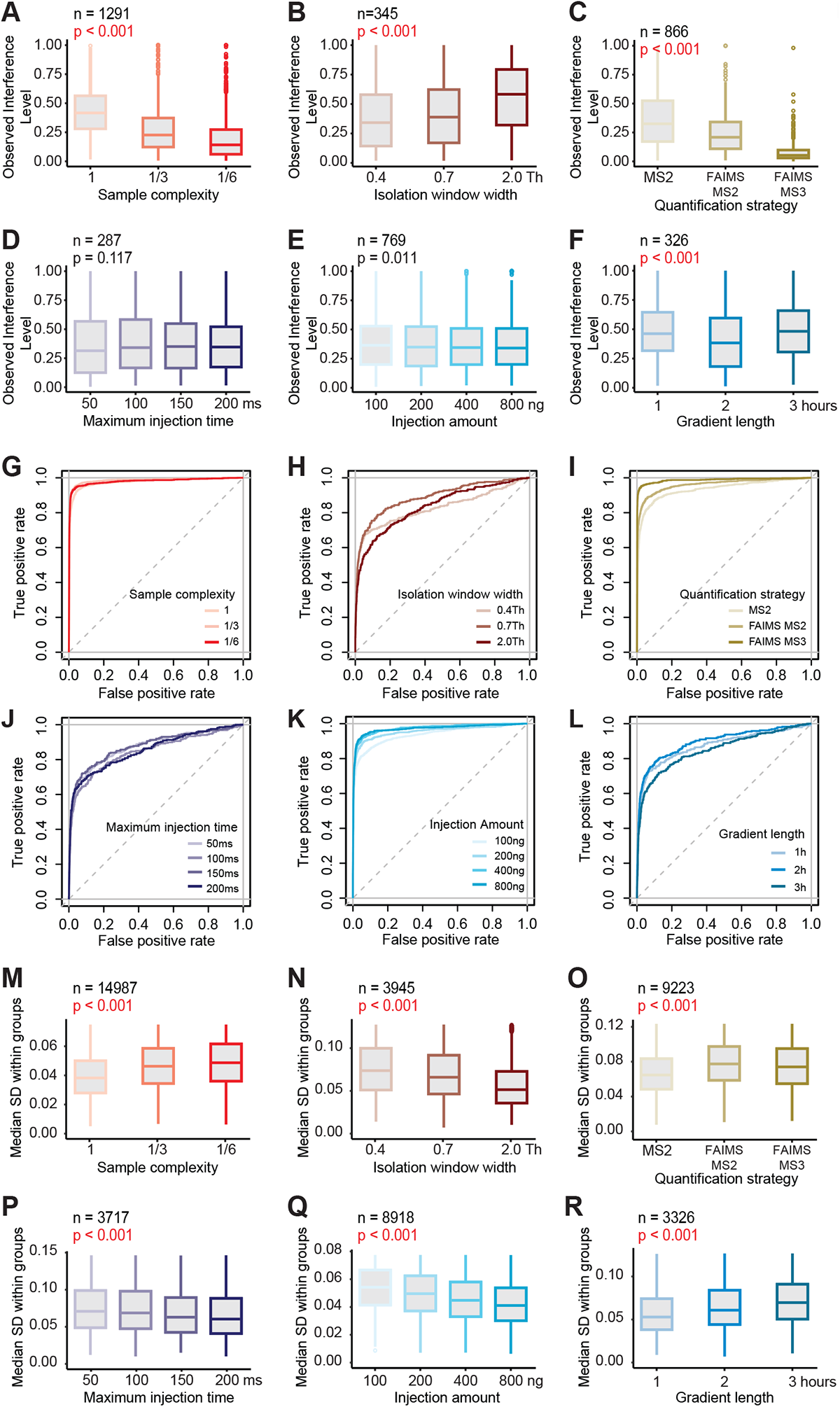
Dependence of ion interference and differential expression analysis on measurement parameters. **(A-F)** Distributions of the observed interference level (OIL) of yeast peptide features independently quantified in all conditions within a comparison. P-values denote statistical significance for overall differences between conditions (Friedman test). The examined measurement parameters are: (A) Varying sample complexities. Numbers 1, 1/3 and 1/6 correspond to factors of sample complexity reduction via fractionation – e.g. 1 implies just one measurement run of the unfractionated sample, while 1/6 implies six measurement runs in total of the six fractionated parts with reduced sample complexity. (B) Varying isolation window widths, in Thomson (Th). (C) Varying quantification strategies. FAIMS-MS3 measurements were also real time-searched (RTS). (D) Varying maximum injection times, in milliseconds (ms). (E) Varying injection amounts, in nanogram (ng). (F) Varying gradient lengths, in hours (h). **(G-L)** ROC-curves calculated from yeast and human peptide features independently quantified in all conditions within comparisons. Measurements are the same as in (A-F). The true positive rate was calculated as the fraction of yeast peptide features correctly classified as differentially expressed at a certain significance level. The false positive rate was calculated as the fraction of human peptide features incorrectly classified as differentially expressed at a certain significance level. **(M-R)** Distributions of the median within-groupstandard deviation of yeast and human peptide features independently quantified in all conditions within a comparison. Measurements are the same as in (A-F). P-values denote statistical significance for overall differences between conditions (Friedman test).

Acetyl (K) and phospho (STY)-peptide enriched samples were measured as follows: LC-MS/MS analysis was performed on an UltiMate 3000 RSLC nano-HPLC System (Thermo Scientific), containing both a trapping column for peptide concentration (PepMap C18, 5 x 0.3 mm, 5 µm particle size) and an analytical column (PepMap C18, 500 x 0.075 mm, 2 µm particle size (Thermo Scientific), coupled to a Q Exactive HF-X Orbitrap (with HCD, higher-energy collisional dissociation mode) mass spectrometer via a Proxeon nanospray flex ion source (all Thermo Scientific). For peptide chromatography the concentration of organic solvent (acetonitrile) was increased linearly over 2 h from 1.6% to 28% in 0.1% formic acid at a flow rate of 230 nL/min. For acquisition of MS2 spectra the instrument was operated in a data-dependent mode with dynamic exclusion enabled. The scan sequence began with an Orbitrap MS1 spectrum with the following parameters: resolution 120 k, scan range 375 – 1,400 m/z, target AGC 3e6, and maximum IT 60 ms. The top 15 precursors were selected for MS2 analysis (HCD) with the following parameters: resolution 45 k, AGC 2e5, maximum IT 200 ms, isolation window 0.7 m/z, scan range 350 – 1500 m/z, and normalized collision energy (NCE) 35. The minimum AGC target was set at 8 × 10^3^, which corresponds to a 4 × 10^4^ intensity threshold. Peptide match was set to preferred. In addition, unassigned, singly and > 6+ charged species and isotopes were excluded from MS2 analysis and dynamic exclusion was set to 30 sec.

### Database search

Raw files were searched with MaxQuant^32^ (version 1.6.14) against a concatenated yeast and human database (Uniprot release 2020.01, canonical sequences) and a common contaminants database, while allowing up to two missed cleavages on tryptic peptides. Search parameters considered oxidation of methionine as well as N-terminal acetylation as variable modifications, and included carbamidomethylation of cysteine residues as a fixed modification. TMTpro label modifications on N-termini and lysine residues were searched as either variable or fixed. Search settings additionally considered variable phospho (STY) or acetyl (K) peptide modifications for raw files of acetyl (K) and phospho (STY)-enriched samples. Raw files generated by FAIMS quantification were split according to scan-specific compensation voltages (CV) using Freestyle (Thermo Fisher). The resulting files were then treated as any other raw file for subsequent database searching. All search results were filtered to reach an expected FDR of 1% at the PSM level.

Some data were generated using alternative search tools **(Table S2)**. *FragPipe*: Thermo raw files were converted to mzML using msConvert^33^ and searched (closed search) against a concatenated yeast and human database including common contaminants using FragPipe^34^ (version 16.0). Carbamidomethylation on cysteines was specified as a fixed; oxidation (methionine), N-terminal acetylation and TMTpro 16plex label were searched as variable. PSM-validation was conducted with Percolator^35^, PSMs were filtered at 1% FDR, and protein inference was conducted with ProteinProphet^36^ filtered at 1% FDR. Additional filters using TMT Integrator: minimum PSM probability: 0.9, minimum best peptide probability: 0.9, normalization: “None”. *Proteome Discoverer*: Raw files were searched against a concatenated yeast/human database using Proteome Discoverer MS Amanda^37^ v2.0. Search parameters: Carbamidomethylation on cysteines was set as static, N-terminal TMTpro 16plex modification as variable modification. Oxidation (methionine), deamidation (asparagine and glutamate), and TMTpro 16plex label were set as dynamic. Maximum number of missed cleavages: 2. Results were filtered at 1% FDR on PSM and protein level using Percolator^35^, and at a minimum MS Amanda Score of 150 at PSM level.

### Raw file data extraction

Apart from using data base search engines, select spectral features like m/z, intensity and noise values of centroided peaks were extracted directly from Thermo raw files via an in-house command line application that uses the Thermo RawFileReader library (https://github.com/fstanek/rawStallion). The combined output was imported into the R statistical environment^38^ and further analyzed there.

### Reporter intensity inference and isotopic impurity correction

Reporter ion intensities were inferred by taking the maximum observed peak intensity value within a ± 0.002 Th window at the expected reporter ion m/z. For this task, we used code from the R proteomics software MSnbase^39^ and utilized the reporter ion classes pre-defined therein. Further, we used linear algebra to correct for TMT isotopic impurities as implemented in MSnbase^39^. The impurity matrix was calculated manually from the product sheet provided by the manufacturer (lot number VC294906).

### Estimation of run-specific peptide density

Retention time and m/z values of all identified, uniquely charged peptide features were used to construct a run-specific two-dimensional kernel density estimate, evaluated on a 200 x 200 grid. The calculation employed the kde2d function from the R package MASS^40^. Resulting density values were rescaled to sum up to 1. Further, the square root of all density values was taken to shrink extreme differences in the m/z and retention time plane, and density values of 0 were substituted with the minimum calculated positive density. Finally, the 200 x 200 density grid was interpolated for each individual PSM at the precursor-specific m/z and retention time coordinates using the interp.surface function from the R package fields^41^.

### Calculation of relevant variables

***OIL:*** The “Observed Interference Level” (OIL) denotes the PSM-specific fraction of interference-induced reporter ion signal at MS2 level. It is calculated as the ratio between the average normalized reporter ion intensity of channels where no quantitative signal is expected (but nonetheless observed due to interference), and the average normalized reporter ion intensity of all other interference-affected channels. To ensure values between 0 and 1, calculated OIL values above 1 are capped at 1. For yeast PSMs in the yeast-human mixture experiment the exact calculation is thus given by

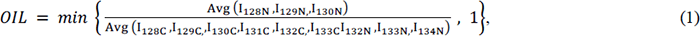

where *I_c_*, *c* ∈ {126*C*, 127*N*, …, 134*N*} is the normalized reporter ion intensity of channel c.

***PPF:*** The “Precursor Purity Fraction” (PPF) denotes the PSM-specific fraction of signal originating from precursor peptide ions (including isotopes) with respect to the total measured signal in the isolation window of bordering MS1 scans. It is similar to existing ion purity metrics like PIF^42^. First, all centroided peak intensities *I_ip_*, *I* ∈ {−1, +1}, *p* ∈ {1, 2, …, *k_i_* } found in the isolation windows of the two bordering MS1 scans indexed by *i*, *i* ∈ {−1, +1} are extracted, where *k*_*_ denotes the total number of distinct peaks in the isolation window of MS1-scan *i*. It is then checked if the precursor ion peak can be found among the peaks in the preceding MS1 scan (*i* = −1) within a margin of error of ± 0.0025 Th. If the check is unsuccessful, the information of the second last MS1 scan is used instead, provided that the precursor ion peak is found there (this was typically the case). We observed this situation to occur more frequently for measurements on Exploris and Eclipse instruments than on HF-X Orbitrap instruments. If the precursor ion peak is still not found, its intensity is imputed at the declared precursor m/z by taking the minimum intensity value over all observed peaks in the preceding MS1 scan (*i* = −1). Concerning the following MS1 scan (*i* = +1), if the respective isolation window is entirely empty, the scan is skipped and not used in the calculation. Else, if the isolation window is non-empty but the precursor ion peak is not found at the declared precursor m/z (± 0.0025 Th), the precursor peak intensity is again imputed as just described. In the rare event that the precursor peak cannot be found in either of the bordering MS1 scans, the following MS1 scan (*i* = +1) is also skipped. Next, the extracted peak intensities *I_ip_*, *i* ∈ {−1, +1}, *p* ∈ {1, 2, …, *k_i_*} in the isolation windows of bordering MS1 scans are inversely weighted by absolute differences between MS1 retention times *t_i_* and the retention time of the MS2-scan *t_Ms2_* that produced the PSM. This results in weighted intensities 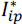:

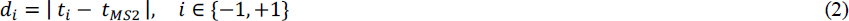

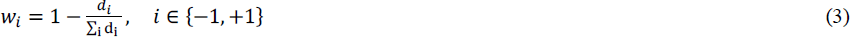

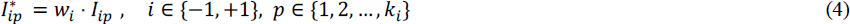

Next, heavy (+1 Da) and light (-1 Da) isotope peaks of the precursor peptide are determined using a mass error tolerance of ± 0.00125 Th. This results in the subset of peaks 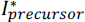 comprising all weighted peak intensities 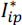 that can be ascribed to the precursor peptide, including heavy and light isotope ions, detected in the isolation window of bordering MS1 scans. The precursor purity fraction PPF is then defined as

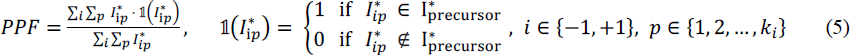

***TIW:*** The “Total Intensity in the Isolation Window” (TIW) is defined as

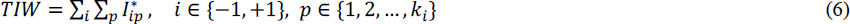

This equation uses the same annotation as in PPF calculation. TIW is the denominator in equation (5).

***PIC:*** The “Total Peptide Ion Current” (PIC) is calculated as the total ion current (TIC) of an MS2 scan minus the total reporter ion intensity of the same scan. This metric therefore equals the sum of all peak intensities stemming from peptide fragment ions in MS2 scans.

### Multiple linear regression modeling

We employed multiple linear regression modeling to model and predict the observable interference-induced reporter ion signal in the yeast-human mixture experiment. The model formula is given by

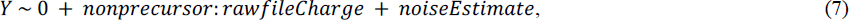

where:

*Y* is the dependent variable of the model and denotes the total reporter ion intensity measured in channels 128N, 129N and 130N of yeast PSMs (i.e. the human-derived reporter ion interference).

*nonprecursor = TIW* ⋅ (1 − *PPF*). This predictor variable therefore denotes the total intensity of non-precursor interfering ions observed in the isolation window.

*rawfileCharge* is a categorical variable describing the charge state of the precursor ion as reported by the mass spectrometer and documented in the raw file. Charge states above 3 were set to 3 to ensure sufficient numbers of observations in each category.

*noiseEstimate* is a PSM-wise estimate of the signal of non-observable interfering ions in the isolation window. It is calculated as the product of the average reported noise value of peaks in the isolation window of the preceding MS1 scan, and the interpolated run-specific peptide density at the precursor-specific m/z and retention time coordinate.

Further, “0” denotes an intercept fixed at 0, and the colon denotes interaction effects between predictor variables. The model parameter coefficients were estimated via robust linear regression modeling with bisquare weighting using the rlm function from the R package MASS^40^. This robust estimation procedure accounts for outliers as well as extreme heteroscedasticity. Data from each unique raw file was modeled with its own separate set of parameters.

Moreover, we propose an extended model that comprehensively explains the total reporter ion intensity of MS2-scans in any MS2-quantified multiplex proteomics experiment. The model formula is given by

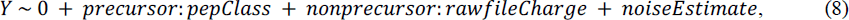

where:

*Y* is the dependent variable of the model and denotes the total reporter ion intensity in MS2-scans.

*precursor = PIC ⋅ PPF*.

*pepClass* is a categorial variable that specifies empirical classes of specific combinations of observed precursor peptide characteristics. It is calculated using a greedy classification algorithm that continuously creates binary splits in peptide characteristic predictor variables in order to best explain observed differences in the fragmentation efficiency of precursor peptides.

*nonprecursor* = *PIC* ⋅ (1 − PPF).

*rawfileCharge and noiseEstimate* are defined as above.

As before, parameter estimates were calculated via robust linear regression modeling with bisquare weighting. Data from each unique raw file was modeled with its own separate set of parameters. Further, separate modeling was applied to data recorded with different compensation voltages in FAIMS measurements.

### Calculation of the Estimated Interference Level (EIL)

Fitting a linear regression model (Equation 8) to MS2-quantified multiplex proteomics data results in estimated model parameters 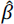 that can be used to predict PSM-wise levels of reporter ion interference in MS2-scans. We define the “Estimated Interference Level” or EIL in short as

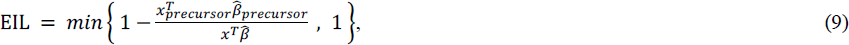

where:

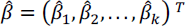 is the k x 1 vector of estimated parameter coefficients,

*x* is a k x 1 vector containing the PSM-specific realizations of predictor variables,

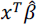 is therefore the fitted value of the model, i.e. the PSM-specific model estimate of total reporter ion intensity.

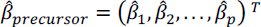 is a p x 1, p < k subset of 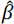 containing only the coefficients that describe the contribution of the precursor peptide to *Y*,

*x_precursor_* is the corresponding p x 1, p < k subset of *x*, such that

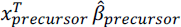 is the PSM-specific model estimate of reporter ion intensity generated only by precursor peptide ions, including heavy and light isotopes.

### Interference correction

Similar to Savitski *et al*.^23^, we propose a PSM-wise interference correction algorithm that generates interference-corrected reporter ion intensities from the estimated interference at MS2 level. The major difference to the original approach lies in using a different kind of impurity metric for the calculation – the estimated interference level (EIL) – instead of an impurity metric that is derived from the MS1 isolation window purity like 1-PPF. Additionally, our algorithm replaces interference-corrected intensity values that fall below a certain threshold with a spectrum-wise minimum. The calculation for the yeast-human mixture experiment is given by

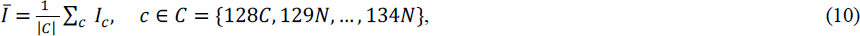

where *I*_c_ denotes the untransformed, normalized reporter ion intensity of reporter channel c; *I̅* thus equals the average normalized reporter ion intensity. Importantly, missing values as well as empty channels are excluded in this calculation. Then, corrected normalized reporter ion intensities 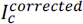 are calculated as

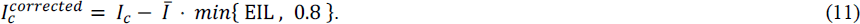

In this formula, EIL values are capped at an arbitrary cutoff of 0.8 to mitigate overcorrection. Finally, corrected reporter intensities 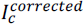 that fall below the minimum of observed MS2 peak intensities are replaced with that minimum to mitigate potential artefacts due to overcorrection. Missing reporter intensity values are substituted equally.

### Data transformation, normalization and general analysis

Reporter ion intensities of were log2-transformed and subsequently normalized using a cyclic LOESS normalization strategy contained in the R package limma^43^ or by using DESeq2’s size factors^44^ for normalization as described previously^45^. After normalization, reporter ion interference was assumed to affect all channels uniformly. Prior to calculating OIL values (Equation 1) and interference-corrected reporter ion intensities (Equation 11), log2-transformed and normalized reporter ion intensities were transformed back from log-space by applying the corresponding inverse function on the transformed data (i.e. exponentiation with base 2). Statistical testing for differential expression between groups was conducted via the Limma-trend testing procedure^46^ on log2-transformed normalized reporter ion intensities. Various additional R packages were used in the analysis and data visualization^47–50^.

## RESULTS

### Experimental design

To interrogate ion interference, we designed an artificial two-proteome multiplex proteomics experiment comprising TMTpro 16plex-labeled tryptic peptides of budding yeast (W303-1A) and human cells (Jurkat E6-1) (**Figure 1A**). Importantly, this system was designed to allow direct observation of reporter ion interference at the MS2 level via the three reporter channels 128N, 129N and 130N in quantified yeast peptides (**Figure 1B**). The profile of total measured yeast and human reporter intensities supports this premise (**Figure 1C**) by illustrating how human signal interfered in yeast peptide measurements. Notably, the mass ratio of human to yeast peptide material was set to approximately 10:1 (**Figure 1A**), thereby ensuring a sufficiently large number of quantified yeast peptides to support the analysis while minimizing the interference coming from yeast. Moreover, to determine the contribution of chemical label background to ion interference, one sample of excessive TMT (channel 126C) label was used without peptides. However, the corresponding signal was negligible in comparison to the interference-induced reporter ion signal in channels 128N, 129N and 130N (**Figure 1C**). This implicates that ion interference is generally not caused by quenched or hydrolyzed TMT label molecules but instead by labeled peptide ions that are co-isolated and co-fragmented during MS2-based quantification of target peptides.

### Measuring interference at MS2 level

Using this direct quantitative readout on interference at MS2 level, we first explored how commonly varying sample and measurement parameters affect interference. In particular we were interested in the factors which, all other things being constant, altered the extent of reporter ion interference at the MS2 level. The six examined parameters comprised the sample complexity (represented by varying degrees of sample fractionation), isolation window width, quantification method (MS2, FAIMS-MS2 and FAIMS-MS3 RTS (real-time search)), maximum MS2 injection time, sample injection amount and HPLC gradient length. To avoid bias, our analysis focused on peptide features with distinct combinations of amino acid sequence, chemical modifications, and charge state. These unique peptide features were then filtered for independent quantification in all measurements to be compared, thereby ensuring that the results are not influenced by varying depths of quantification or the stochasticity inherent to data-dependent acquisition (DDA). For each yeast peptide feature, we determined the level of interference at MS2 level by calculating the average fraction of interference-induced reporter ion intensity (i.e. intensity in the yeast-free channels 128N, 129N, 130N) to the reporter ion intensity across all other channels. This fraction was termed the “Observed Interference Level” (OIL) (**Equation 1**). It would assume 0 in the absence of any measurable reporter ion interference in channels 128N, 129N and 130N, and could reach up to 1, at which point the entire quantitative signal in a yeast scan was attributed to human-derived interference. Our findings support previous literature results^7–10, 51^ by showing that reducing sample complexity (**Figure 2A**) or isolation window widths (**Figure 2B**) effectively alleviate interference at the MS2 level. We also observed a previously described reduction in interference for FAIMS-MS2 quantification over standard MS2 quantification^13^, and almost completely interference-free TMT quantification via FAIMS-MS3 RTS^10^ (**Figure 2C**). Varying maximum MS2 injection times (**Figure 2D**), as well as varying sample injection amounts (**Figure 2E**) yielded no remarkable change in reporter ion interference. However, our data show lowest levels of interference for intermediate gradient lengths (**Figure 2F**). We attribute this finding to altered peptide elution profiles, which – combined with the applied instrument settings – result in the fragmentation of peptides at more favorable elution times^52^. Notably, all the above-mentioned changes in interference are equally reflected by an accompanied degree of ratio compression in calculated yeast peptide fold changes (**Figure S1A-F**).

Next, we investigated how these changes in ion interference impact differential expression (DE) analysis. This is of particular interest since ratio compression evidently reduces the effect size of group differences (**Figure S1A-F**), which in turn negatively impacts statistical power. Reporter intensity values were first log2-transformed to ensure normality of errors. We then conducted pairwise differential expression testing via the Limma-trend testing procedure^46^ between the two sample groups 100:9 (channels 131C, 132C, 133C) and 100:6 (channels 128C, 129C, 130C), resulting in the theoretical fold changes of 1.0 for human and 1.5 for yeast peptides, respectively (**Figure 1A**). ROC curves were calculated to compare the diagnostic abilities of correctly classifying yeast peptide features as differentially expressed versus human peptides as non-differentially expressed. The data revealed that ion interference is not the only factor affecting DE analysis results in isobaric labeling-based quantification, since the varying levels of interference (**Figure 2A-F**) failed to sufficiently explain ROC curve differences (**Figure 2G-L**). In addition, the reporter ion signal strength appears pivotal by influencing measurement precision (i.e. variance within groups) via an observable mean-variance trend (**Figure S2**), which in turn affects statistical power. We observed significant changes in reporter ion signal (**Figure S1G-L**) and corresponding within-group variances (**Figure 2M-R**) for all comparative measurements, which ultimately help to explain differences in ROC-curves when paired with changes in observed interference levels (**Figure 2A-F**). For example, despite higher levels of interference in the more complex samples (**Figure 2A**), the increased precision due to the higher overall signal (**Figure 2M** and **Figure S1G**) compensated for the loss of accuracy in statistical testing (**Figure 2G**). In other words, we observed that the presence of additional reporter ion interference signal evoked a tradeoff between accuracy and precision, with both roughly balancing each other out in their combined influence on DE analysis. Importantly, in order to focus on the effect of sample complexity in isolation, we intentionally did not scale up the total injection amount across all samples of reduced complexity compared to the unfractionated sample. Incidentally, an increase in the total injection amount showed improved classification results due to a gain in precision from higher overall reporter ion signal (**Figure 2K**, **2Q** and **Figure S1K**). Hence, performing fractionation combined with an increased total sample load distributed across multiple measurements should benefit DE analysis while at the same time mitigate the effect of ratio compression.

Regarding isolation window widths, our data indicated that the increased precision due to an overall increase in reporter ion signal with larger isolation window widths (**Figure 2N** and **Figure S1H**) can lead to improved classification results (**Figure 2H**), even if that comes at the cost of higher levels of interference and ratio compression (**Figure 2B** and **Figure S1B**). Notably, in addition to a general higher transmission efficiency, window widths of 0.7 Th also manage to co-isolate +1 and -1 isotopes of triply charged precursor peptides, thus providing a possible explanation as to why a window of 0.7 Th classified markedly better than a window of 0.4 Th (**Figure 2H**). On the other hand, a window width of 2.0 Th, with a median OIL above 0.5 (meaning that for more than 50% of quantified yeast peptides at least half of the signal was in fact human-derived) appeared to suffer in accuracy to such an extent that the accompanied increase in precision could not compensate for the loss in statistical power due to waning accuracy (**Figure 2B**, **2H**, **2N** and **Figure S1B**).

Comparisons between ROC-curves of varying maximum injection times and gradient lengths were complicated by differing tendencies to repeatedly quantify identical peptide features due to changes in measurement speed and measurement time, respectively. This somewhat hampers the direct comparability and interpretation of calculated ROC curves. Still, the improved classification result from the measurement with an intermediate gradient length is striking (**Figure 2L**), which emphasizes the importance of reconciling the chromatographic separation with MS instrument settings that determine the quantification of peptides at specific points in the elution profile for maximum accuracy and precision^52^.

Finally, given the applied instrument settings (**Table S1**) we observed FAIMS-MS2 and in particular FAIMS-MS3 RTS quantification to considerably outperform standard MS2 quantification in correctly classifying differential expression (**Figure 2I**). Both FAIMS-MS2 and FAIMS-MS3 quantification seemingly benefitted from increased accuracy due to a notable reduction of interference (**Figure 2C** and **Figure S1C**) without losing too much in precision (**Figure 2O**). Nevertheless, it is important to put these results into a broader context. So far, we focused on unique peptide features that were independently quantified across all compared conditions in order to avoid any selection bias in the analysis. However, variation in the addressed measurement parameters often resulted in substantial differences in the number of unique quantified peptides (**Figure S1M-R**). Altogether, we find that maximizing identifications, fold change accuracy (i.e. minimizing ion interference) and correct classification of differential expression at the same time is not trivial because there is a tradeoff between the three. The choice of measurement strategy is thus best guided by the available instrumentation, as well as the specific aims and constraints of the particular experiment.

### Modeling interference at MS2 level

To gain a more causal and mechanistic understanding of reporter ion interference generation, we examined the observed reporter ion interference of yeast peptide scans in more detail. We speculated that the observed interference level (**Equation 1**) would be best explained by the total ion composition in the precursor peptide’s isolation window range, which gives direct insight into the degree of unwanted co-isolation and subsequent co-fragmentation for MS2-based quantification. This reasoning is not novel. For example, the “Precursor Ion Fraction”^42^ (PIF), implemented in the quantitative proteomics software platform MaxQuant^32^, and other similar ion purity metrics^34, 37^ estimate the degree of precursor purity by calculating the fraction of ion signal attributed to the precursor peptide with respect to the total ion signal in the isolation window of bordering MS1 scans. Usually, these purity metrics are used to filter out PSMs considered too impure for reliable quantification. Here, we introduce a purity metric termed “Precursor Purity Fraction” (PPF) which equally ranges from 0 to 1 by definition (**Equation 2-5**). Calculated PPF values correlate well with existing purity metrics (**Figure S3A-C**), but perform best in the modeling approach described below due to subtle differences in calculation (see methods).

Remarkably, regardless of which purity metric is considered, the MS1 isolation window purity appears as a rather weak predictor for the actual level of interference at MS2 level (**Figure 3A**). In fact, interference at the MS2 level is substantially underestimated, and even PSMs that appear pure in their respective isolation windows (i.e. PPF = 1) exhibited considerable reporter ion interference (**Figure S4**). We attribute this discrepancy to how Orbitrap raw files are recorded during measurement. To minimize raw data file size, signals below a certain threshold are considered “noise” and consequently removed. What remains as reduced information in Orbitrap raw spectra are thus a) all peaks with intensities above calculated local noise thresholds that serve as a cutoff; and b) noise values specific to each centroided peak (**Figure 3B**), which are proportional to the aforementioned local noise thresholds^f1^. We assume that by removal of these low-intensity signals, a substantial part of interfering ions remains hidden in the MS1 isolation window range, especially when signal to noise ratios of target precursor ions are low. At the same time, we hypothesize that the noise values contained in the raw data could still sufficiently represent the potential strength of the hidden signal. To explore these hypotheses, we employed multiple linear regression modeling. The dependent variable of the model was defined as the spectrum-wise total measured interference at MS2 level, calculated as the summed reporter ion intensity of channels 128N, 129N and 130N of yeast peptide measurements. In line with our previous observation (**Figure 3A)**, a model that only accounts for the observable portion of ion interference in the MS1 isolation window range performed poorly in predicting the reporter ion interference at MS2 level (**Figure 3C**, upper). Next, we adapted the model formula by including a regressor variable that aimed to reflect the potential noise contribution. Assuming that co-isolated interfering ions hidden within the spectral noise are in fact other labeled peptide ions of relatively low abundance, we calculated this additional regressor variable by combining two numerical sources of information via multiplication: First, a measure that reflects the potential maximum intensity of such ions, calculated as the average noise value in the precursor peptide’s isolation window range; and second, the estimated frequency of such ions, taken from a measurement-specific empirical peptide density that spans the m/z-retention time plane (**Figure 3D**). The updated model (**Equation 7**) managed to accurately predict the interference-induced reporter ion signal for all yeast peptide measurements (**Figure 3C**, lower), including PSMs that appeared pure in their respective MS1 isolation window range.

**Figure 3.**
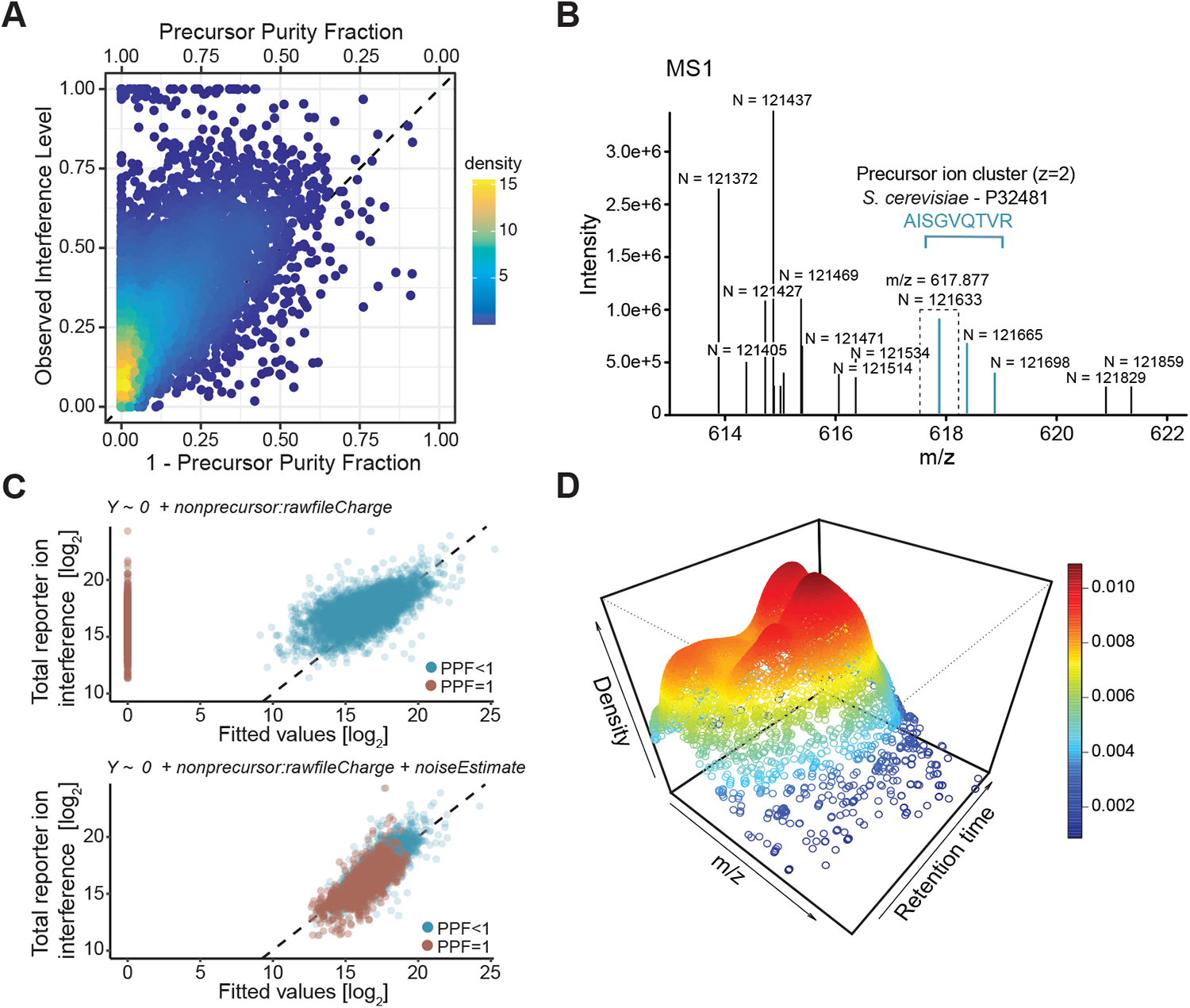
Modeling reporter ion interference. **(A)** Relationship between observed interference levels (OIL) and MS1 isolation window impurities (1-PPF). Each data point corresponds to a single yeast PSM. **(B)** Example of a recorded MS1 spectrum (raw file “20201030_[…]_complexity_P1”, scan number 12583) in which peaks are annotated with their respective noise values N. The turquoise peak at m/z=617.877 corresponds to a yeast precursor peptide ion that was subsequently targeted for MS2-based quantification at an isolation window width of 0.7 Th (indicated by dashed lines). Despite the absence of visible interference at MS1 level (PPF=1), the resulting PSM with MS2 scan number 12589 exhibited substantial reporter ion interference at MS2 level (OIL=0.39). **(C)** Model prediction of the measured reporter ion interference by two nested linear regression models. Model formulas are depicted on top of each plot. Each data point corresponds to a single yeast PSM. The dashed line represents the identity function (y = x) and therefore reflects a perfect prediction. A pseudo-count of 1 was added to each fitted value before log-transformation in order to ensure finite values. **(D)** 3D-density map showing density estimates calculated for all PSMs in a single measurement run (raw file “20201030_[…]_complexity_P1”). A higher estimated density signifies a larger number of unique peptide features quantified in the vicinity of the m/z and retention time coordinate.

### Estimating interference at MS2 level

Accurate estimation of ion interference at MS2 level could prove extremely valuable for the qualitative and quantitative interpretation of MS2-quantified multiplex proteomics experiments. As shown above, a linear regression model fitted to the measured data has the potential to provide this crucial information. However, implementation of such a linear regression model requires direct measurement of reporter ion interference at MS2 level to support model training, which is usually not the case in biological experiments.

We therefore adjusted the model to make it applicable to all MS2-quantified multiplex proteomics data. Consequently, the dependent variable was redefined as the total measured reporter ion signal of an MS2 spectrum, containing both interference and non-interference reporter ion signal. Most of this signal is expected to originate from the precursor peptide ion that was targeted for MS2-based quantification and it is essential that the model accurately reflects this contribution to the total signal. However, this task is complicated by a striking divergence in the relationship between precursor and reporter ion intensities, presumably due to peptide-specific differences in the fragmentation efficiencies (**Figure 4A**). A part of this variation could be explained with differences in precursor ion charge state (**Figure 4B**). However, even within individual charge states, a significant level of heterogeneity remained unexplained. Therefore, we tested whether this could be attributed to other peptide characteristics. Remarkably, we found that the majority of the observed variance could be attributed to distinct empirical precursor peptide classes (**Figure 4C** and **4D**). Apart from the precursor ion charge state, these peptide classes were characterized by the number of TMT labels as well as the presence or absence of specific amino acids. Interestingly, we found that empirical peptide classes also varied greatly in frequency and composition between individual data sets, most likely due to sample and measurement-specific variables (e.g. normalized collision energies, chromatography, etc.). Ultimately, we developed a greedy decision tree algorithm that automatically infers an optimal set of empirical peptide classes for all PSMs from a unique measurement run (**Figure 4D**). Class variables determined in this way could then be incorporated as separate categorical regressor variables into the linear regression model.

**Figure 4.**
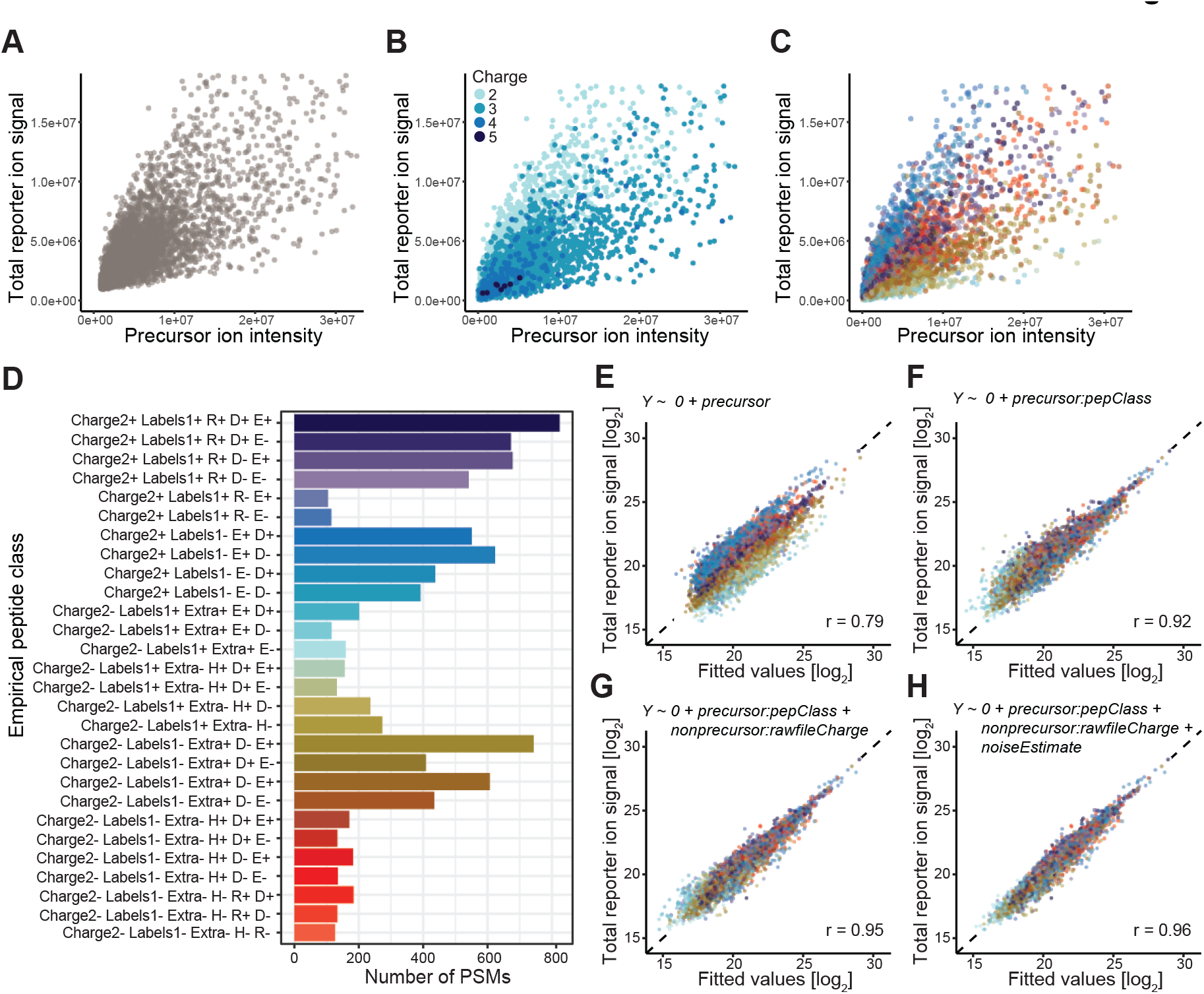
Modeling total reporter ion signal. **(A)** Dependence of total reporter ion signal on precursor ion intensity in the MS1 isolation window range. Each data point corresponds to a single yeast or human PSM from raw file “20201030_[…]_complexity_P1”. **(B)** The same relationship as in (A), colored by precursor ion charge states. **(C)** The same relationship as in (A), colored by empirical peptide classes of varying fragmentation efficiency. **(D)** Overview of empirical peptide classes shown in (B), as determined by a greedy decision tree-based classification algorithm. Peptide class labels reflect paths in the decision tree from root to leaf (i.e. terminal node). Each split (i.e. internal node) represents the action of further dividing a peptide class into two distinct sub classes based on a single physicochemical property (e.g. contains arginine vs. contains no arginine, symbolized by “R+” and “R-”, respectively) such that the resulting within-class variance is minimized. The variables available for creating new splits are: The precursor charge state (Charge); the number of TMT-labels (Labels); the presence and absence of amino acids histidine (H), arginine (R), lysine (K), glutamate (E) and aspartate (D); and the existence of an additional positive charge that is not explained by the total number of charged functional groups in the peptide’s chemical structure (Extra). The algorithm ensures a sufficient number of observations within each class (>100) by preventing splits that reduce the number of PSMs below that threshold. **(E-H)** Model prediction of the measured total reporter ion signal by four nested linear regression models. Model formulas are depicted on top of each plot. Each data point corresponds to a single yeast or human PSM. The dashed line represents the identity function (y = x) and therefore reflects a perfect prediction. An increasing number of predictor variables were sequentially included into the model from top left to bottom right. Note that differences in slopes will appear as differences in y-intercepts since both axis are log-transformed. r denotes the calculated Pearson correlation coefficient.

Further, we encountered a second hurdle. In most datasets, linear model fitting was made difficult by a nonlinear relationship between the total MS1 intensity signal in the isolation window (**Equation 6**) and the total reporter ion signal at MS2 level **(Figure S5A-B**, upper rows). This might be explained by a nonlinear signal response in the Orbitrap. While complex signal processing steps are implemented to address this issue^f1^, a perfect linear relationship between MS2 intensities and their corresponding MS1 precursor intensities might not be achieved in all cases. Consequently, our model had to be adjusted. We substituted the total intensity in the isolation window with the total intensity of peptide fragment ions, which is calculated as the total ion current (TIC) of an MS2 scan minus the recorded total reporter ion intensity of the same scan. We termed this metric “Total Peptide Ion Current” or PIC in short. Notably, the relationship between PIC and the total reporter ion signal demonstrated a robust linear relation **(Figure S5A-B**, lower rows). Nevertheless, MS2 scans also fail to reveal the full extent of low-intensity signals contributing to the noise. Therefore, an updated model based on MS2 information still has to account for interference to faithfully predict the total reporter ion signal. Ultimately, we arrived at a multiple linear regression model formula that comprehensively describes the total reporter ion signal in MS2 scans (**Equation 8**), making it applicable to any MS2-quantified multiplex proteomics experiment.

Fitting this model to the yeast-human PSM data resulted in improved prediction results with increasing numbers of relevant predictor variables (**Figure 4E-H**). Due to the strong predictive performance of the final model (**Figure 4H**), we were able to determine the individual contributions of the model terms to the overall predicted reporter ion signal in the experiment. Independent model fitting (on the six lower complexity samples) revealed that about 20 percent of the total reporter ion signal originates from ion interference (**Figure S6**). Approximately half of this signal could be attributed to noise, i.e. interfering ions that are not directly visible in the MS1 isolation window range.

It is noteworthy that this global interference prediction strategy can also be applied at the PSM level. Therefore, we were able to assess the relative contribution of reporter ion interference to the total reporter ion signal for each specific PSM. These individual estimates were then aggregated at the peptide and protein levels. We coined this model-based prediction of the fraction of reporter ion interference at MS2 level the “Estimated Interference Level” (EIL) (**Equation 9**), which can be thought of as another ion purity metric like the previously introduced precursor purity fraction (PPF). Yet unlike metrics of MS1 isolation window purity, EIL accurately estimates ion interference, with higher precision when aggregating from PSMs to peptides or proteins (**Figure 5A**).

**Figure 5.**
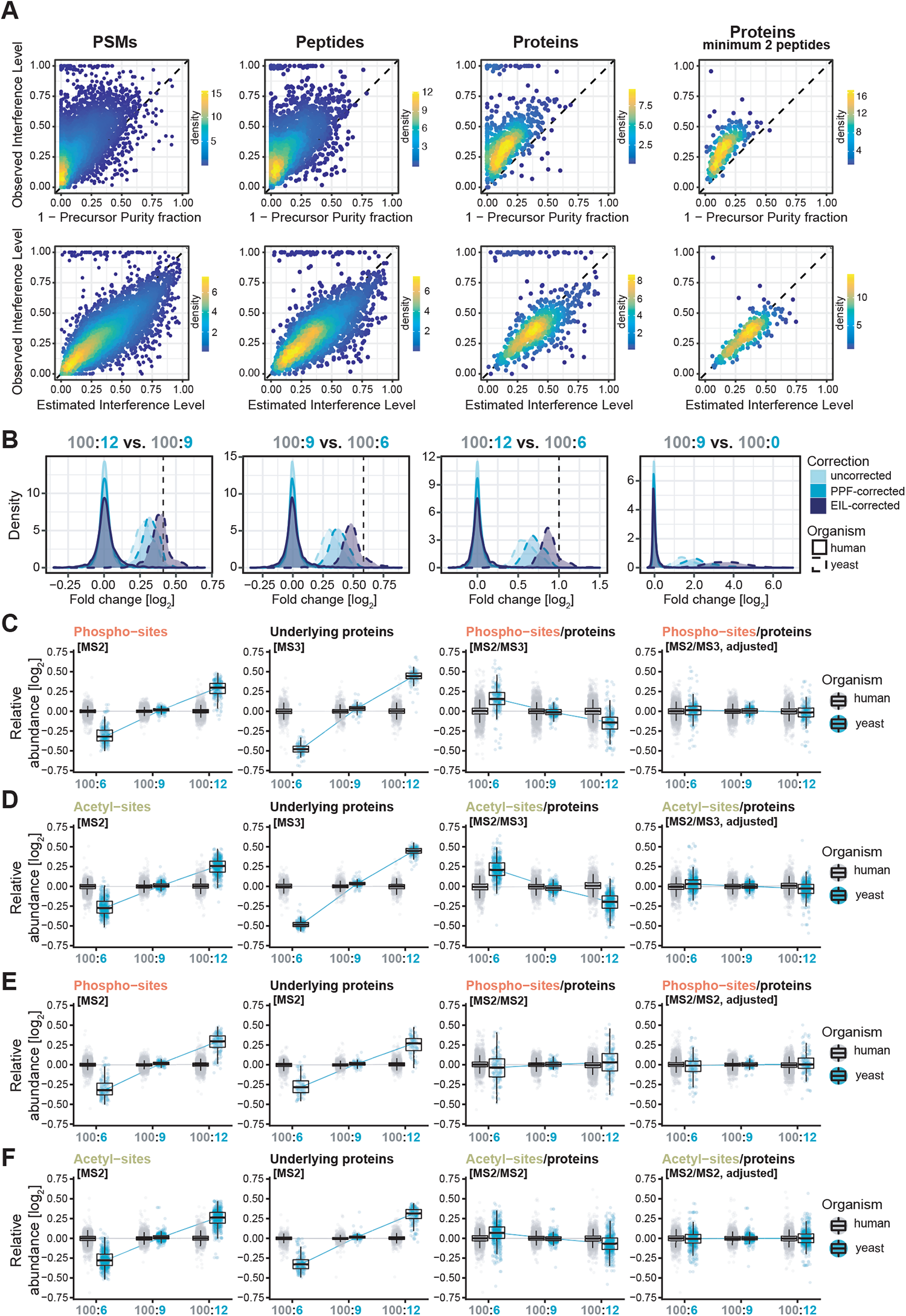
Calculating and utilizing accurate interference estimates. **(A)** Relationship between observed interference levels (OIL) and MS1 isolation window impurities (1-PPF) (top row), or estimated interference levels (EIL) (bottom row). Columns reflect distinct levels of yeast feature aggregation (PSMs, peptides, proteins, and proteins filtered for at least 2 unique and unambiguous peptides), with increasing levels of aggregation from left to right. OIL values were calculated based on feature-wise reporter ion intensities (aggregated by summation). EIL and PPF values were aggregated by calculating weighted averages of PSM-specific values, with weights reflecting the intensity contribution of individual PSMs to the aggregated reporter ion intensity. **(B)** Distribution of log2-transformed fold changes of yeast and human proteins, resulting from the pairwise group comparisons 100:12 vs. 100:9, 100:9 vs. 100:6, 100:12 vs. 100:6 and 100:9 vs. 100:0 (left to right). The dashed black line in each plot marks the expected theoretical fold change for yeast proteins and constitutes 1.33, 1.5, 2.0 and infinite, respectively. All proteins were additionally filtered for at least 2 unique and unambiguous peptides to reduce the potential for misspecifications of human peptides as yeast. **(C-F)** Results of site-to-protein normalization. Rows correspond to independent normalization results of phospho- or acetyl-sites, normalized to either MS3-quantified proteins (C-D) or MS2-quantified proteins (E-F). Columns show data before and after normalization. Each data point corresponds to a single yeast or human feature’s relative abundance in the respective group after log2-transformation and group-averaging.

To validate the EIL-based estimation of interference, we applied our modeling strategy to other datasets. First, we selected published datasets^53, 54^ of MS2-quantified TKO9 and TKO11 proteomic standards^53^. These commercial TMT-based standards each comprise three multiplexed congenic yeast knockout (KO) strain proteomes which allow direct observation of interference at MS2 level for peptides of the missing proteins. Paolo *et al*.^53^ took advantage of this property to calculate the so-called “Interference-Free Index” (IFI), which translates to 1-OIL in our study. We compared OIL to model-based EIL values of the three knocked-out proteins, and found a much-improved prediction of reporter ion interference over conventional purity metrics (**Figure S7A-B**). Second, we applied the EIL-based modeling procedure to a MS2 dataset we recorded with FAIMS. Since FAIMS only partially reduces interference (**Figure 2C** and **Figure S7C**, upper panel) we interrogated whether EIL-based modeling can improve interference removal. Indeed, we observed strongly improved prediction of interference (**Figure S7C**, lower panel). Finally, we tested our approach on lower complexity samples with an expected lower level of interference. For this purpose, we used samples enriched for lysine-acetylated or phosphorylated (STY) peptides generated from the yeast-human ground-truth experiment. Again, the results emphasized the robustness and accuracy of EIL-based prediction of reporter ion interference, and the advantage over conventional purity metrics, in these samples (**Figure S7D-E**).

### Interference correction

Equipped with accurate estimates of reporter ion interference, we aimed to effectively correct ratio compression in any dataset. Savitski *et al*. calculated interference-corrected reporter ion intensities using a conventional MS1-based purity estimate (termed “Signal-to-Interference” or S2I)^23^. We adapted this approach by utilizing model-based EIL values instead, along with minor changes that aim to mitigate potential over-correction (**Equation 10-11**). We evaluated our algorithm’s performance based on theoretical fold changes of yeast-derived proteins of 1.33, 1.5, 2.0, and infinite in our yeast-human mixture experiment (**Figure 1A**). Conversely, human proteins had uniform relative abundances across all samples, thus mimicking proteins of unchanged expression levels. We compared three different modes of correction: No interference correction; correction based on the conventional MS1 isolation window purity metric PPF; and correction based on the estimated interference level EIL. The results highlight several key aspects. First, there is a notable improvement compared to the original correction approach (**Figure 5B**). Second, EIL-based interference correction considerably decreased – but did not completely remove – ratio compression (**Figure 5B**). Regarding this second point, we speculate that EIL-based interference correction has the potential to completely decompress fold changes – after all, EIL values manage to accurately estimate interference at MS2 level (**Figure 5A** and **Figure S7**). However, in the dataset presented here this might have been prevented by a faint background of yeast peptide-derived interference. Since different amounts of yeast peptides were used in our experiment (**Figure 1**) such an interference contribution would be *non-uniform*. This could have biased the between-sample normalization, ultimately diminishing group differences. Additionally, due to the applied ceiling of 0.8 for EIL during interference correction (**Equation 11**), a slight degree of ratio compression might have remained. Third, while accuracy increased, the correction procedure resulted in a loss in precision (**Figure 5B**). This was especially true for the human protein population, which exhibited an increased spread of log2 fold changes after correction. Notably, this tradeoff between accuracy and precision matches the already observed tradeoff that a change in interference entails (**Figure 2** and **Figure S1**), yet here it is induced entirely computationally. Differential expression and ROC-analyses further illustrate that, despite drastic differences in quantitative accuracy, the classification results remain relatively unchanged (**Figure S8**).

### PTM site-to-protein normalization

Multiplex proteomics is particularly well-suited for the study of post-translational modifications, thanks to the joint processing of samples in one pool and the low number of missing values per labeling set. However, quantitative data on the PTM level is inherently confounded by the overall changes in underlying protein abundance^55^. In order to accurately interpret results, differential abundances in modified peptides must be calibrated to protein levels, which can be assessed by quantifying unmodified peptides of the same proteins. This approach enables inference of changes on site level that are independent of changes on protein level, reflecting alterations in the post-translational modification status^55, 56^. Ideally, a simple normalization strategy, such as dividing site intensities by corresponding protein intensities or site fold changes by corresponding protein fold changes, can reveal this crucial information. However, in multiplexed proteomics this operation is complicated by ion interference, which compresses relative group differences on site and protein level independently. This introduces a risk for bias in site-to-protein normalized abundances or site-to-protein normalized fold changes, potentially leading to an incorrect reflection of biology.

Using our artificial yeast-human samples as a system with known ground-truth, we investigated this problem in more detail. MS2-quantified yeast and human phospho- and acetyl-site intensities were calibrated to both MS2 as well as FAIMS-MS3 RTS quantified protein signatures by arithmetic division (**Figure 5C-F**, column 1-3). In theory, existing group differences in yeast site abundances should level out after normalization to yeast proteins known to exhibit the same relative abundance pattern. Instead, the calculated ratios suggested varying acetylation and phosphorylation rates between groups (**Figure 5C-F**, column 3). For example, yeast sites calibrated to MS3-quantified proteins revealed a trend that is the exact opposite of the original yeast abundance pattern (**Figure 5C-D**, column 3). This highlights the danger of over-calibration when normalizing interference-affected site intensities to MS3-quantified protein intensities with little to no interference. On the other hand, calibrating to MS2-quantified proteins produced ratios that were more accurate on average, but displayed large variation (**Figure 5E-F**, column 3). This observation underscores that similar overall levels of ratio compression are not sufficient to ensure unbiased normalization results, as individual site and protein pairs can still differ in their respective interference levels.

In light of these results, there is a need for unbiased site-to-protein normalization. Since the bias directly stems from unequal degrees of interference in individual site and protein pairs, we reasoned that adjusting for this disparity prior to ratio-building would remove this bias. We thus leveraged our model-based EIL values to establish such an algorithm. In brief, for each site and protein feature pair, reporter ion interference was artificially added to the feature with lower EIL until reaching equal levels. This adjustment only minimally affects the quantitative data before ratio-building, and further does not increase within-group variances, as opposed to the interference correction approach described above. Following this adjustment to reach equal interference levels, normalization was performed as before via division. The resulting site-to-protein normalized abundances managed to accurately reflect the expected absence of between-group changes in site-to-protein ratios (**Figure 5C-F**, column 4). Moreover, the ratios displayed a much-reduced spread, particularly when normalized to MS2-quantified proteins (**Figure 5E-F**, column 4). Given these results, we think that this revised normalization approach promises effective reduction of bias from ion interference, which clears the way for the discovery of real biological changes at the post-translational level through isobaric labeling-based quantification.

## DISCUSSION

Despite extensive research into reporter ion interference in multiplex proteomics^5–12, 22–29, 52–54, 57–60^, a comprehensive mechanistic explanation of this phenomenon remained difficult. We attribute this challenge to two primary factors. Firstly, our study reveals that a significant portion of the signal associated with ion interference is not directly observable due to its removal during signal processing in Orbitrap instruments. Secondly, several factors influencing interference need to be considered when trying to understand, quantify, and mitigate interference. Consequently, previous efforts to correct ratio-compressed TMT data have primarily employed a phenomenological approach, relying on a single measured variable to gauge the extent of ratio compression^6, 8, 9, 23–25, 52^ . However, since these single variables are not sufficient to fully account for the actual interference, the effectiveness of proposed computational methods has been limited. In contrast, technical solutions such as MS3 methods or charge state reduction^12–14, 16^ have demonstrated more promising outcomes. Nevertheless, these alternatives rely on specific hardware requirements and introduce trade-offs concerning acquisition speed and signal strength. In our study, we aimed to address ratio compression by developing a holistic understanding, combining various approaches to determine the underlying causes of ion interference. We utilized a specially designed experimental system to directly observe interference-induced reporter ion signals and ultimately employed an exploratory computational model to identify and quantify the factors contributing to interference.

Our findings revealed that, on average, approximately 20% of the reporter ion signal was attributed to interference, a range similar to the findings reported by Wenger *et al*. who reported around 32% using a wider isolation window^12^. These results highlight the difficulty of achieving accurate relative quantification in multiplex proteomics experiments and emphasize the need for appropriate correction measures. In addition, we investigated the influence of various parameters on the observed interference level and how they eventually affect differential expression testing. These parameters included sample complexity, isolation window width, quantification method, maximum injection time, injection amount, and HPLC gradient length, and the results from this extensive screening are consistent with previously published reports^6–10, 13, 14, 16, 25, 52, 53^. However, our analysis reveals that these parameters cannot be considered separately, since there are trade-offs between identification rate, quantitative accuracy and precision, and statistical power in DE analysis. Hence, multi-parameter optimization to reach high identification rates, accurate fold change measurements, and correct classification of differentially expressed entities at the same time is a complex task. Our findings provide a guideline for selecting and optimizing the most relevant parameters in TMT multiplexing experiments.

The main goal of this study, however, was to perform a comprehensive characterization of interference to determine its cause and ultimately correct it with the use of computational models. Our analyses clearly show that interference is caused by TMT-labeled peptides or their putative fragments. Quenched TMT-label or its side-products are negligible. Moreover, our modeling approach shows that peptide ion parameters such as charge, amino acid composition, and number of bound TMT labels are important determinants of interference that need to be considered in interference correction. A key finding, however, was that the major contribution to interference is mainly caused by ions that are hidden in the noise of the spectrum. As a consequence, estimates of precursor purity based on the MS1 spectrum, such as PIF or similar^32, 34, 37, 42^, are poor measures of interference and not well suited for interference correction. It was therefore necessary to introduce a novel metric for estimating ion interference, the model-based estimated interference level or EIL. Whether the “hidden” interference derives mainly from labeled peptides or from peptide fragments produced during ionization, as suggested by Erickson *et al*.^61^ cannot be answered from our data. Most likely both factors contribute, although previous observations suggesting that interfering ions are mostly singly charged indicate that the contribution of fragments might be substantial^12^. This would also fit our observation that the interference signal must be rather low and uniform to be fully removed during “de-noising” in the course of spectrum processing.

Building on our comprehensive understanding of the underlying mechanisms of interference, we adjusted our computational model to ensure broad applicability across all TMT datasets. A major consequence of this strategy was the computability of the EIL metric, which provides an accurate estimate of interference at PSM level. Our analysis convincingly demonstrated the effectiveness of EIL in correcting for ratio compression on a global scale. With our approach, we achieved higher quantitative accuracy with similar statistical power in differential expression analysis when compared to analyses using uncorrected data. As a result, we developed a software tool that enables correction of ratio compression that can be applied to any MS2-quantified TMT dataset^62^.

We further benchmarked our interference estimation strategy by testing its effectiveness for PTM datasets. Calibrating relative abundances of modified peptides to the according protein level is crucial for the assessment of changes in PTM patterns under different experimental conditions. However, PTM and protein information suffer from different degrees of compression, which often results in quantification artifacts when simply dividing site intensities by corresponding protein intensities. Our approach offers a solution to this problem, and we demonstrate its effectiveness in two scenarios: when PTM and protein information are both determined using MS2 measurements, and when PTM information is obtained through MS2 while protein information is obtained through MS3 measurements. We have made all the required code available as documented repositories on GitHub to enable the community to apply our method^62, 63^.

Finally, there are important assumptions and limitations that need to be considered. Firstly, we assumed that the isolation efficiency is uniform across the whole isolation window, even though it is non-uniform^64^ and even varies between instruments. However, given the relatively small deviation from uniformity, we do not anticipate a significant impact on our modeling approach. Alternatively, one could experimentally determine the isolation efficiency for a given instrument or infer it from the data to refine the modeling. Secondly, we assumed that every ion peak at MS1 level corresponds to a labeled peptide ion, disregarding the possibility of singly charged, non-labeled, non-peptide ion contamination. To address this, one could computationally remove singly charged signals or use fractionation/FAIMS to remove contamination if necessary. Furthermore, isobaric labeling experiments are mostly performed on Orbitrap FT-mass spectrometers due to the required instrument resolution for multiplexing. However, in the future, other mass spectrometers, such as TOF instruments, might be used that differ in signal detection and processing methods^65^, which could affect ion interference. Thus, the modeling strategy will need to be adapted accordingly. Additionally, ion mobility separation technologies other than FAIMS, such as the trapped ion mobility cell in timsTOF (Bruker), may remove ion interference and provide more accurate data^26^. Another limitation that needs to be considered is the assumption that most proteins in the experiment have similar expression levels, leading to interference affecting all reporter channels evenly. While this assumption is generally valid, it may not always hold true. Therefore, when significant differences exist between multiplexed samples, an alternative correction strategy involving the filtering and/or differential weighting of individual proteins’ PSMs based on measurement accuracy confidence, such as ion purity metrics calculated from the isolation window, may be more appropriate. Such strategies have been previously discussed and implemented by others^23, 27, 57^ and the EIL metric presented in this study can enhance their performance. However, it is important to note that strict filtering of PSMs will always result in a loss of identification numbers.

In summary, our study offers valuable insights into the nature of ion interference, as well as guidance on minimizing interference and optimizing DE testing in multiplex proteomics data. We believe that our work enhances the overall understanding of this fundamental problem and provides new computational strategies for effectively coping with ion interference and ratio compression.

## ABBREVIATIONS

AGC: Automatic gain control
DDA: Data dependent acquisition
DE: Differential expression
EIL: Estimated interference level
FAIMS: High-Field Asymmetric-Waveform Ion Mobility MS
ID: Identification
IFI: Interference free index
IT: Injection time
LFQ: Label free quantification
MS: Mass spectrometry
MS2: MS/MS
MS3: MS/MS/MS
OIL: Observed interference level
PIC: Total peptide ion current
PIF: Precursor ion fraction
PPF: Precursor purity fraction
PSM: Peptide spectrum match
PTM: Post-translational modification
ROC: Receiver operating characteristic
RTS: Real-time search
TIC: Total ion current
TIW: Total intensity in isolation window
TMT: Tandem mass tag

## ACKNOWLEDGEMENTS

We thank Teresa Preglej and Wilfried Ellmeier for providing Jurkat cells, Sabrina Jenull and Karl Kuchler for providing yeast cells used in setup experiments, and Gerhard Dürnberger for the support with the Proteome Discoverer software platform. We are grateful to David Hollenstein and Johannes Griss for their critical feedback on the manuscript, and Andreas Bögehold and Thermo Fisher Global Support for the correspondence on Orbitrap measurement details. We thank the Vienna Biocenter Core Facilities (VBCF) for providing the LC-MS instrument pool. MM, NH, WR and MH were supported by the Austrian Science Fund (FWF) Special Research Program F70.

## AUTHORS’ CONTRIBUTION

**Moritz Madern**: Conceptualization, Data curation, Formal analysis, Investigation, Methodology, Project administration, Software, Validation, Visualization, Roles/Writing – original draft, Writing – review & editing. **Wolfgang Reiter**: Supervision, Visualization, Roles/Writing – original draft, Writing – review & editing. **Florian Stanek**: Software. **Natascha Hartl**: Investigation. **Karl Mechtler**: Resources. **Markus Hartl**: Conceptualization, Funding acquisition, Project administration, Resources, Supervision, Roles/Writing – original draft, Writing – review & editing. All authors approved the final manuscript.

## DATA AVAILABILITY

The mass spectrometry proteomics data have been deposited t o the ProteomeXchange Consortium via the PRIDE^66^ partner repository with the dataset identifier PXD040449. A detailed summary of all measurement raw files and their contribution to the individual results of this study is listed in **Table S2**. The code used in the analysis is available on GitHub^67^. Additionally, we have created two documented GitHub repositories that provide the computational and statistical methods presented in this study in form of ready-to-use R scripts^62, 63^.

## SUPPLEMENTAL DATA

This article contains supplemental data.

## UNPUBLISHED OBSERVATIONS AND PERSONAL COMMUNICATIONS

f1 Footnote 1: personal communication with Thermo Fisher Scientific.

## SUPPLEMENTAL FIGURE LEGENDS

**Figure S1.**
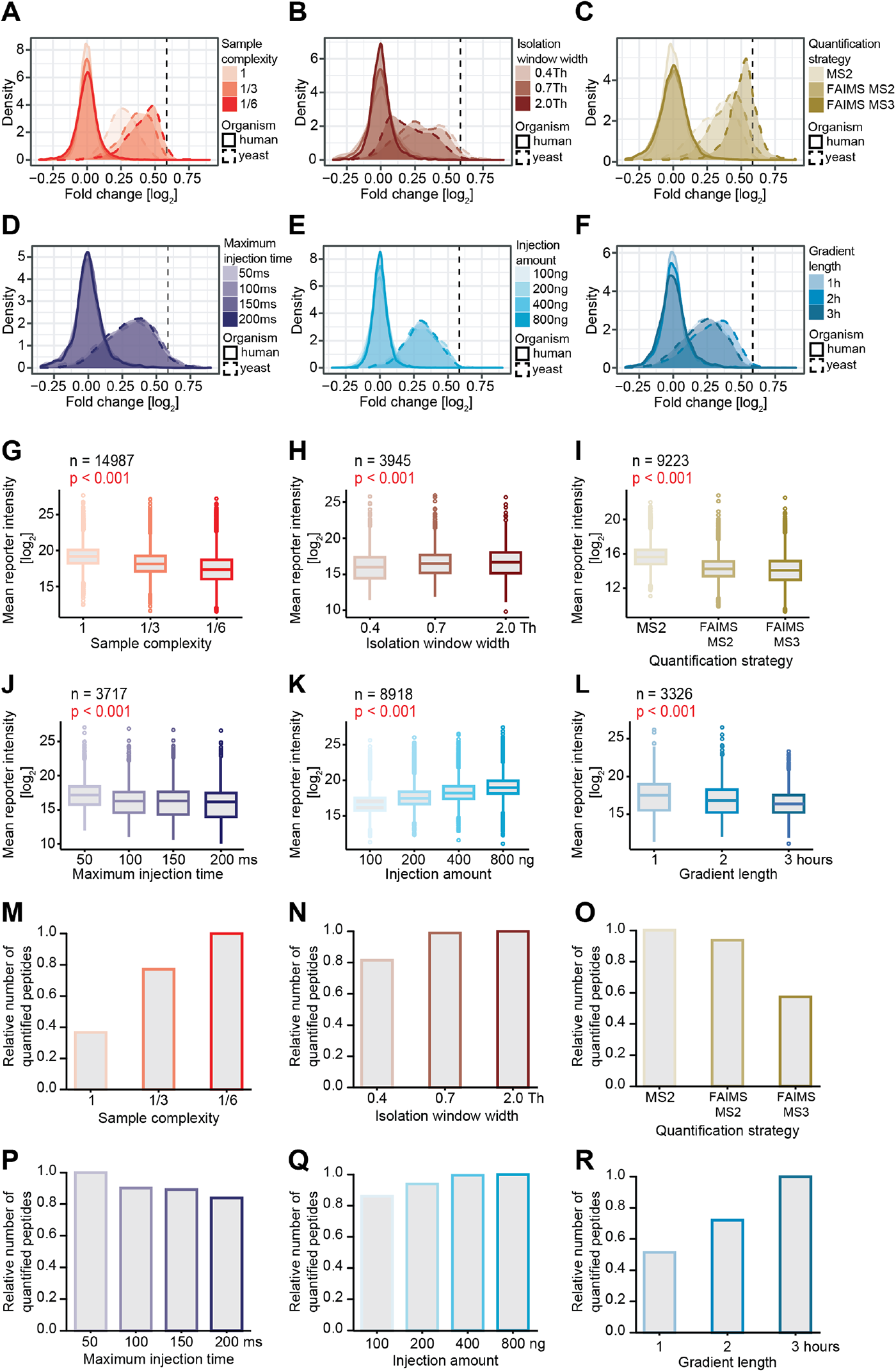
**Supplement to Figure 2. (A-F)** Distribution of log2-transformed fold changes of yeast and human peptide features independently quantified in all conditions within a comparison, resulting from the comparison between groups 100:9 and 100:6. The dashed black line marks the theoretical fold change for yeast peptides. **(G-L)** Distribution of average reporter intensities of yeast and human peptide features independently quantified in all conditions within a comparison. P-values denote statistical significance for overall differences between conditions (Friedman test). **(M-R)** Relative number of unique peptides quantified per condition. The highest number in each comparison was scaled to 1.

**Figure S2.**
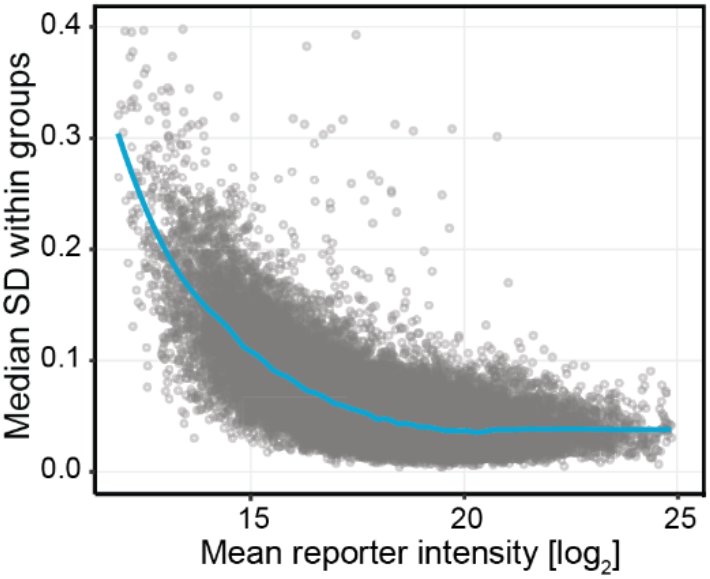
Mean-variance trend in isobaric labeling-based quantification. Empirical mean-variance trend of log2-transformed reporter ion intensities of MS2-quantified yeast and human peptides. The turquoise line represents a loess-fit with a span parameter of 0.05. Each data point corresponds to a single PSM.

**Figure S3.**
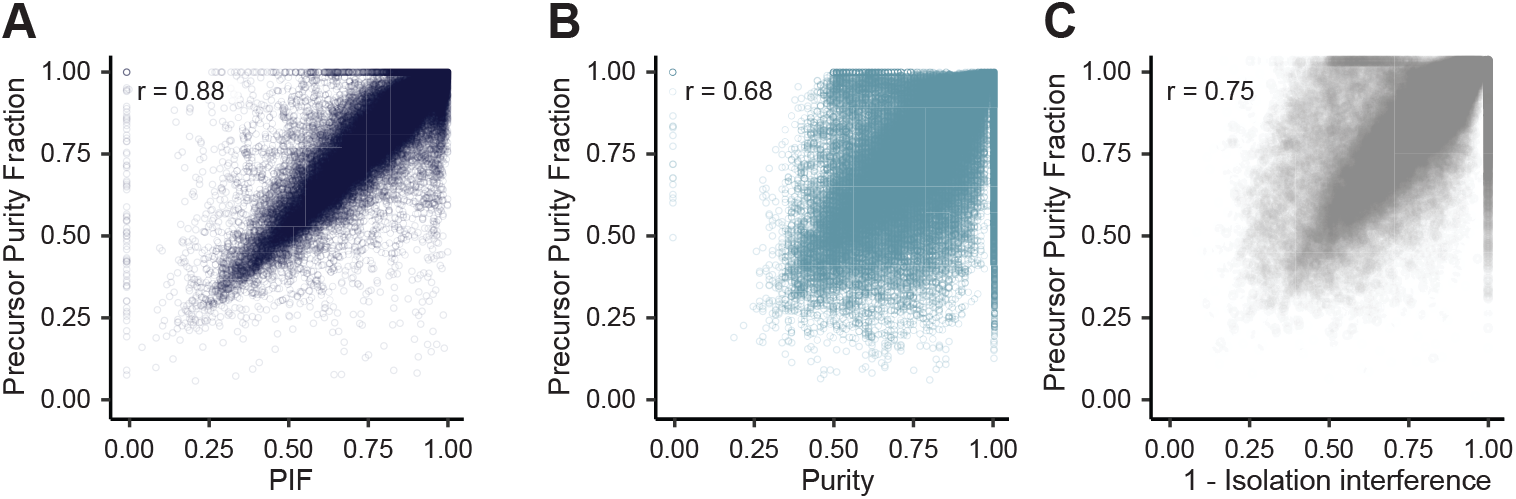
Comparison of Precursor Purity Fraction (PPF) to existing purity metrics. Each data point corresponds to a single PSM. r denotes the calculated Pearson correlation coefficient. **(A)** Comparison with the “PIF” metric implemented in MaxQuant. **(B)** Comparison with the “Purity” metric implemented in FragPipe. **(C)** Comparison with the “Isolation Interference” metric implemented in the Proteome Discoverer software MS Amanda.

**Figure S4.**
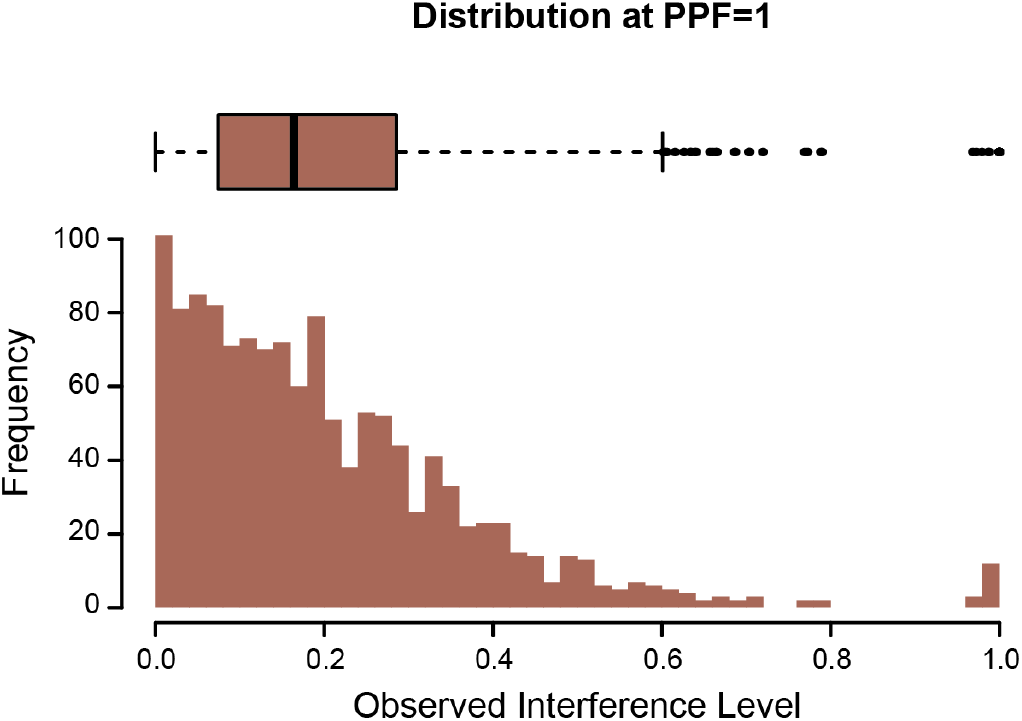
**Supplement to Figure 3A**. Distribution of observed interference levels (OIL) of yeast PSMs at PPF=1. Note that only a small fraction of yeast PSMs actually appears free of reporter ion interference (i.e. OIL=0). The sharp rise in counts at OIL=1 is assumed to stem from human peptides that were misspecified by the search engine as yeast peptides at an FDR of 1% at PSM level.

**Figure S5.**
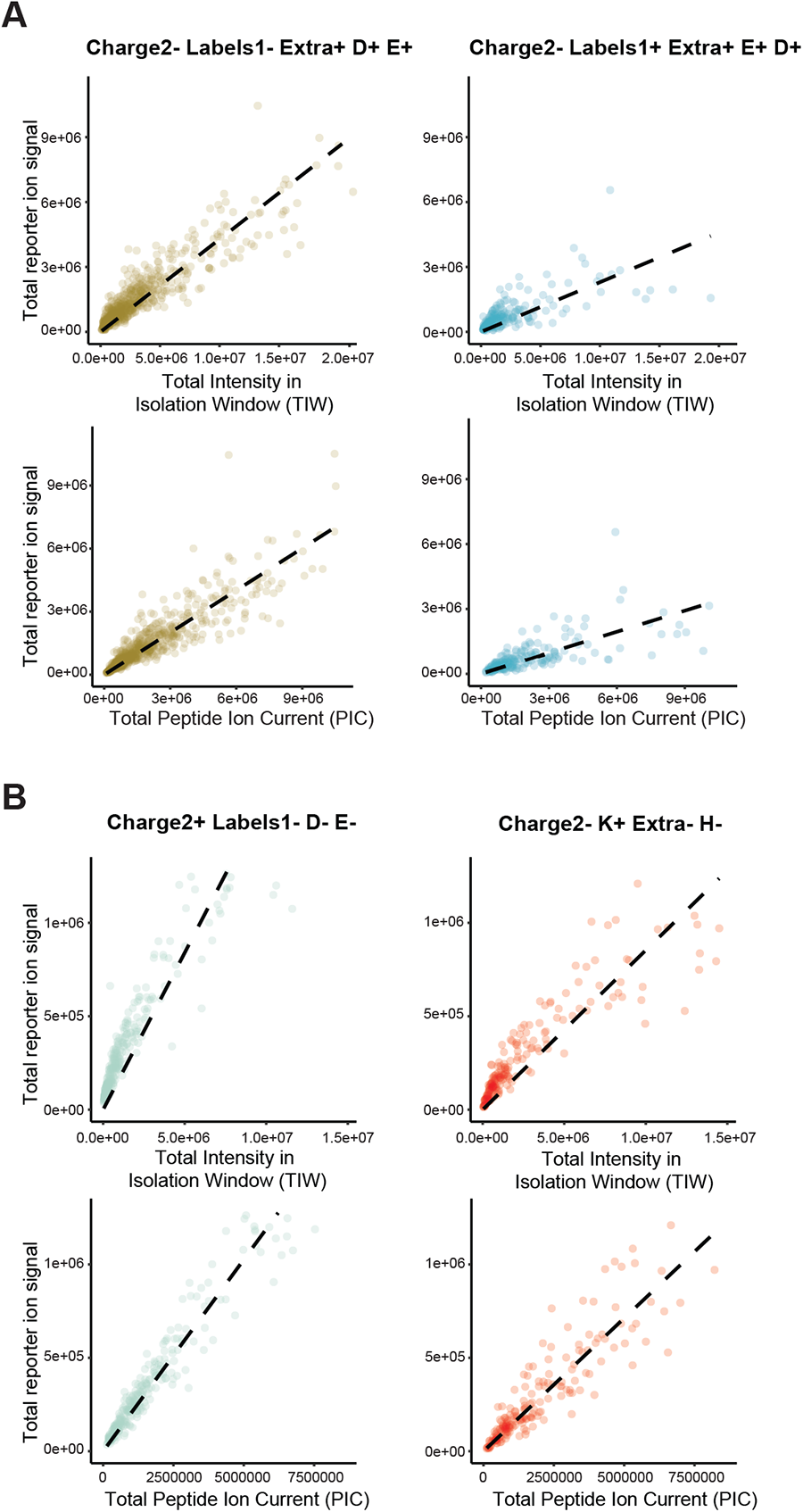
Linearity in regression modeling. Dependence of the total reporter ion signal on the total intensity in the isolation window (TIW) (upper rows) and the total peptide ion current (PIC) (lower rows) for PSMs of two distinct empirical peptide classes per unique dataset. The dashed line in each plot reflects a simple linear regression fit. **(A)** Data of the yeast-human mixture dataset (raw file “20201030_[…]_complexity_P1”). **(B)** Data of a TKO9 dataset published with the original publication (PMID: 27400695).

**Figure S6.**
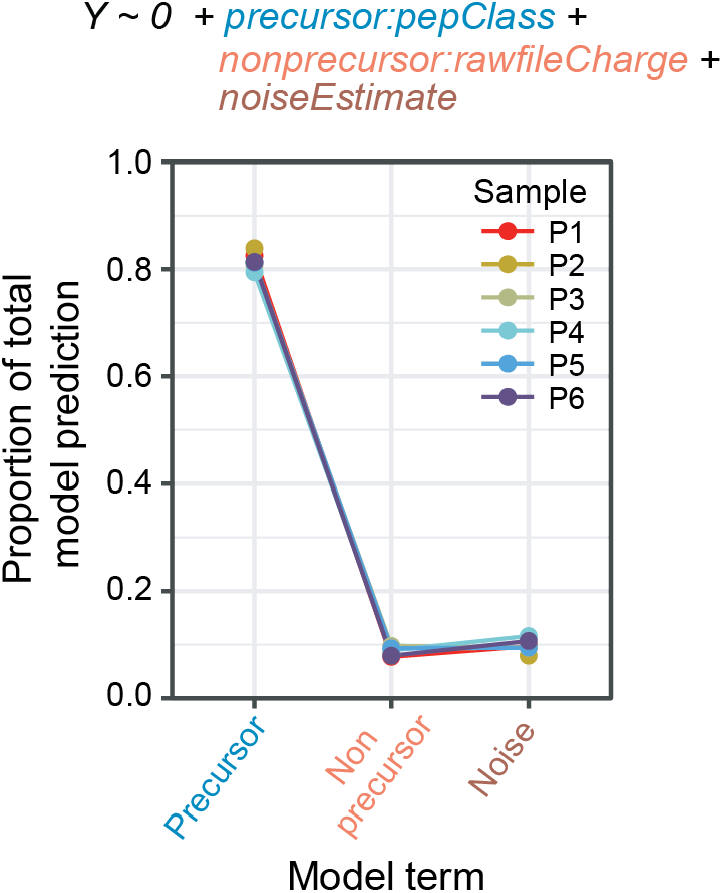
Partition of model prediction. Results display model term-specific proportions of predicted reporter ion signal with respect to total predicted reporter ion signal across all PSMs of a single measurement run. The model was fit separately for each of the six samples of reduced sample complexity (P1-P6) measured via MS2-based quantification. The three model terms (x-axis) relate to different parts of the model equation, as coded by color. The second term (Non precursor) and the third term (Noise) denote the visible and the invisible contribution of ion interference, respectively, to the total reporter ion signal at MS2 level.

**Figure S7.**
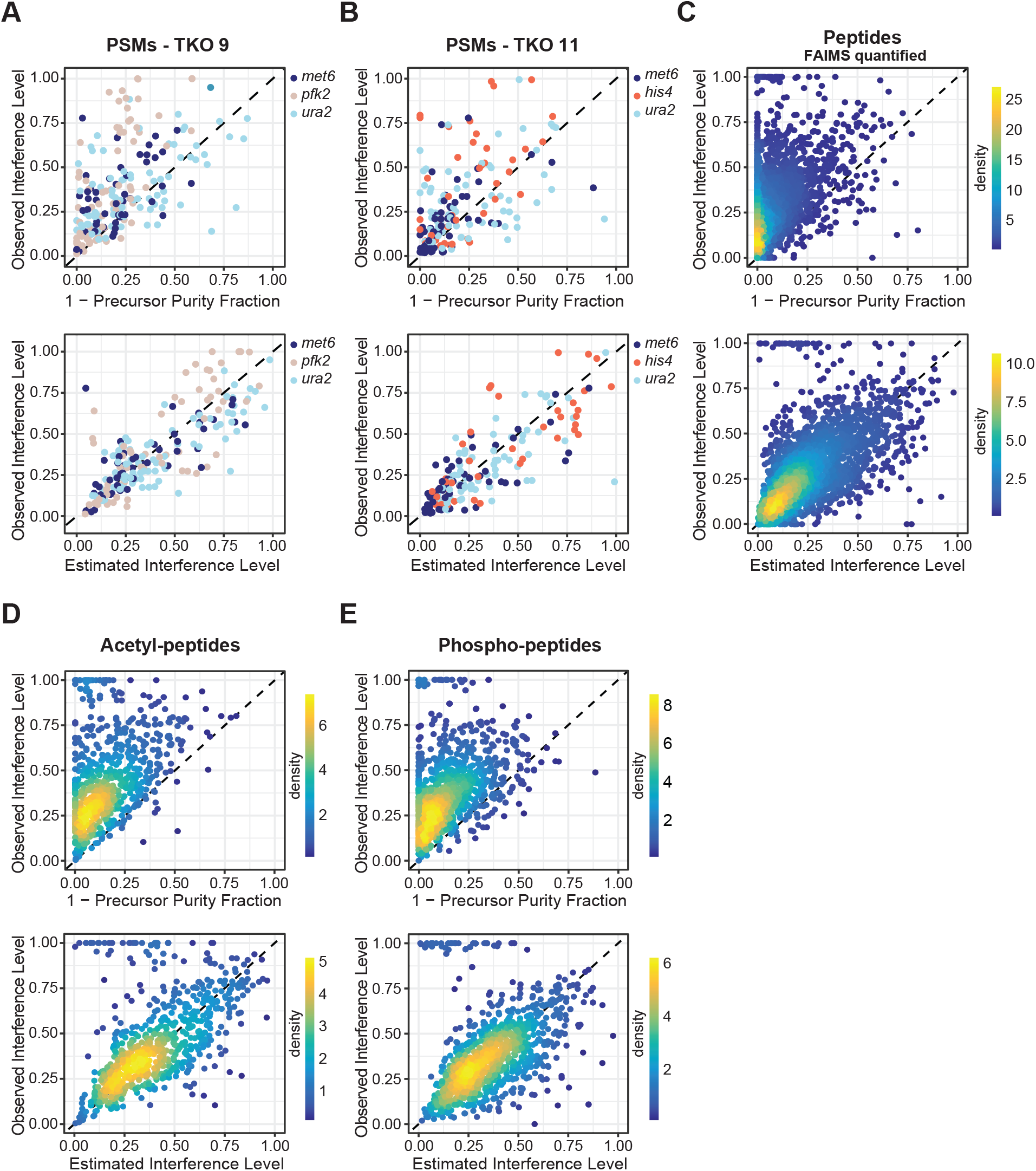
Evaluation of EIL in other data sets. Plots show the relationship between observed interference levels (OIL) and MS1 ion impurities (1-PPF) (top row) or estimated interference levels (EIL) (bottom row). **(A)** PSM level data of previously published triplicate measurements of the TKO9 proteomics standard (PMID: 27400695). Observed interference levels (OIL) were determined for quantified peptides of the three KO genes met6, pfk2 and ura2. **(B)** PSM level data of previously published triplicate measurements of the TKO11 proteomics standard (PMID: 32202424). Observed interference levels were determined for quantified peptides of the three KO genes met6, his4 and ura2. **(C)** Peptide level data of the yeast-human mixture experiment using FAIMS-MS2 quantification. Observed interference levels (OIL) were determined for yeast peptides. Measurements are the same as shown in Figure 2C (FAIMS-MS2). **(D)** Acetyl-peptide level data of the yeast-human mixture experiment using MS2 quantification. Observed interference levels (OIL) were determined for acetylated yeast peptides. **(E)** Phospho-peptide level data of the yeast-human mixture experiment using MS2 quantification. Observed interference levels (OIL) were determined for phosphorylated yeast peptides.

**Figure S8.**
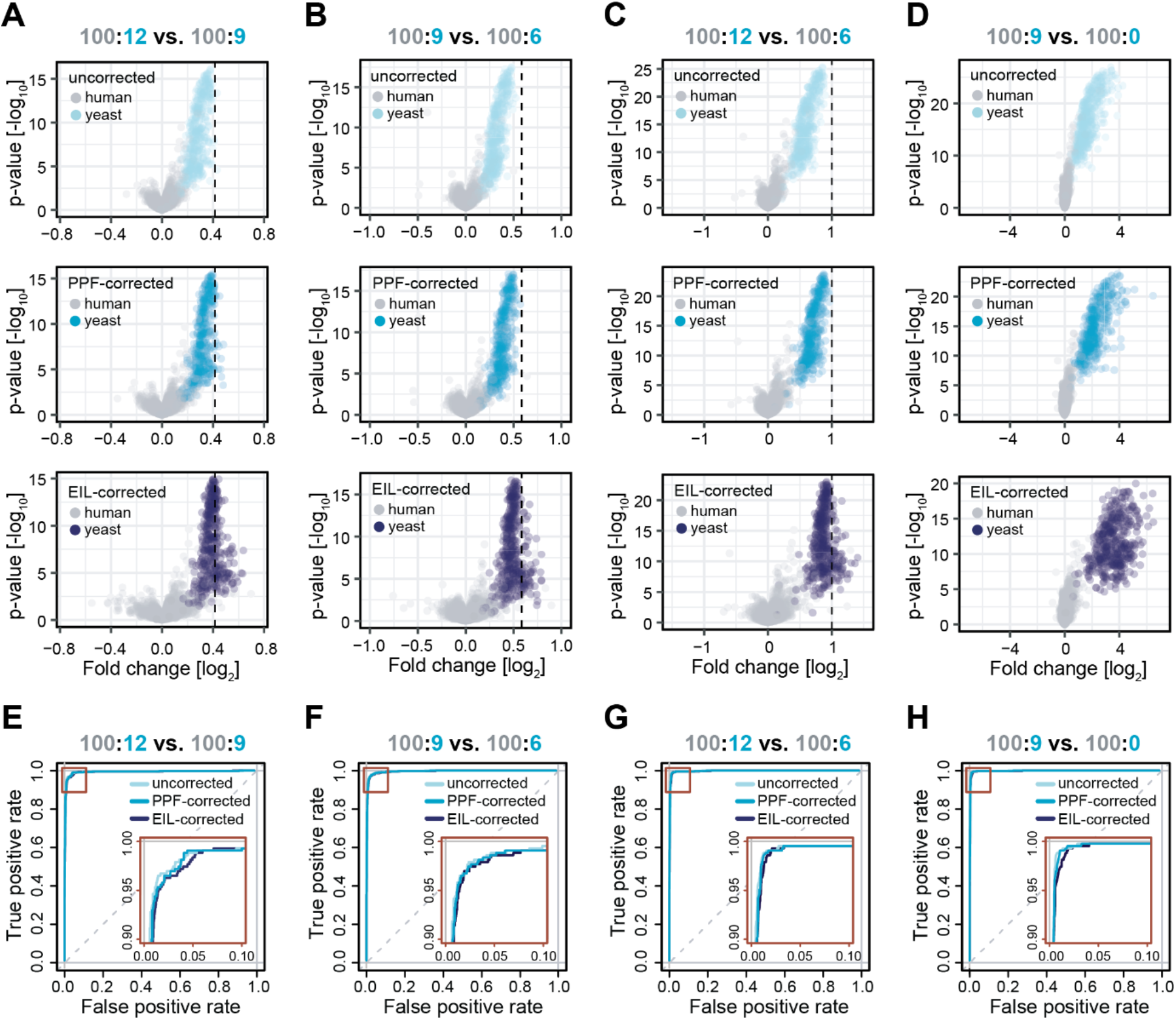
**Supplement to figure 5B. (A-D)** Volcano plots illustrating results of protein level DE testing and fold change accuracy and precision for three distinct interference correction strategies. Columns represent the different pairwise group comparisons already shown in Fig 5B: (A) 100:12 vs. 100:9; (B) 100:9 vs. 100:6; (C) 100:12 vs. 100:6; and (D) 100:9 vs. 100:0. The dashed black line in each plot marks the theoretical fold change for yeast proteins and constitutes (A) 1.33, (B) 1.5, (C) 2.0 and (D) infinite, respectively. Rows represent different interference corrections strategies: uncorrected, PPF-corrected and EIL-corrected (top to bottom). Each data point corresponds to a single yeast or human protein. All proteins were additionally filtered for at least 2 unique and unambiguous peptides to minimize potential misspecification of human as yeast and vice-versa. Statistical testing was performed via the Limma-trend testing procedure on log2-transformed protein intensities. Aggregation to protein intensities was performed by summation of individual PSM’s normalized and interference-corrected reporter ion intensities. **(E-H)** ROC-analyses based on the DE testing results. The true positive rate was calculated as the fraction of yeast proteins correctly classified as differentially expressed at a certain significance level. The false positive rate was calculated as the fraction of human proteins incorrectly classified as differentially expressed at a certain significance level.

**Table S1.** Supplement to Figure 2C, 2I and 2O. Summary of instrument parameters for comparative measurements using different quantification strategies. All measurements were performed on the same instrument (Eclipse) to ensure comparability.

| General measurement settings |  |  |  |  |  |  |  |  |  |  |
| --- | --- | --- | --- | --- | --- | --- | --- | --- | --- | --- |
|  | CVs | MS instrument | Number of samples | Gradient length | Cycle time | MS1 res | MS1 AGC target | MS1 maximum IT | Isolation window | Intensity threshold |
| MS2 | - | Eclipse | 2 | 2h | 3s | 120k | 400000 | 50ms | 0.7Th | 25000 |
| FAIMS-MS2 | -40, -55, -70 | Eclipse | 2 | 2h | 1.2s | 60k | 400000 | 50ms | 0.7Th | 25000 |
| FAIMS-MS3 RTS | -40, -55, -70 | Eclipse | 2 | 2h | 1.2s | 120k | 400000 | 50ms | 0.7Th | 10000 |

| MS2 scan settings |  |  |  |  |  | MS3 scan settings |  |  |  |  |  |
| --- | --- | --- | --- | --- | --- | --- | --- | --- | --- | --- | --- |
|  | MS2 detector type | MS2 AGC target | MS2 collision energy | MS2 Orbitrap resolution | MS2 maximum IT | MS3 detector type | MS3 AGC target | MS3 collision energy | MS3 Orbitrap resolution | MS3 maximum IT | MS3 Number of notches |
| MS2 | Orbitrap | 150000 | HCD 34% | 50k | 100ms | - | - | - | - | - | - |
| FAIMS-MS2 | Orbitrap | 150000 | HCD 34% | 50k | 100ms | - | - | - | - | - | - |
| FAIMS-MS3 RTS | IonTrap | 10000 | CID 30% | - | 50ms | Orbitrap | 150000 | HCD 50% | 50k | 150ms | 10 |

**Table S2.**
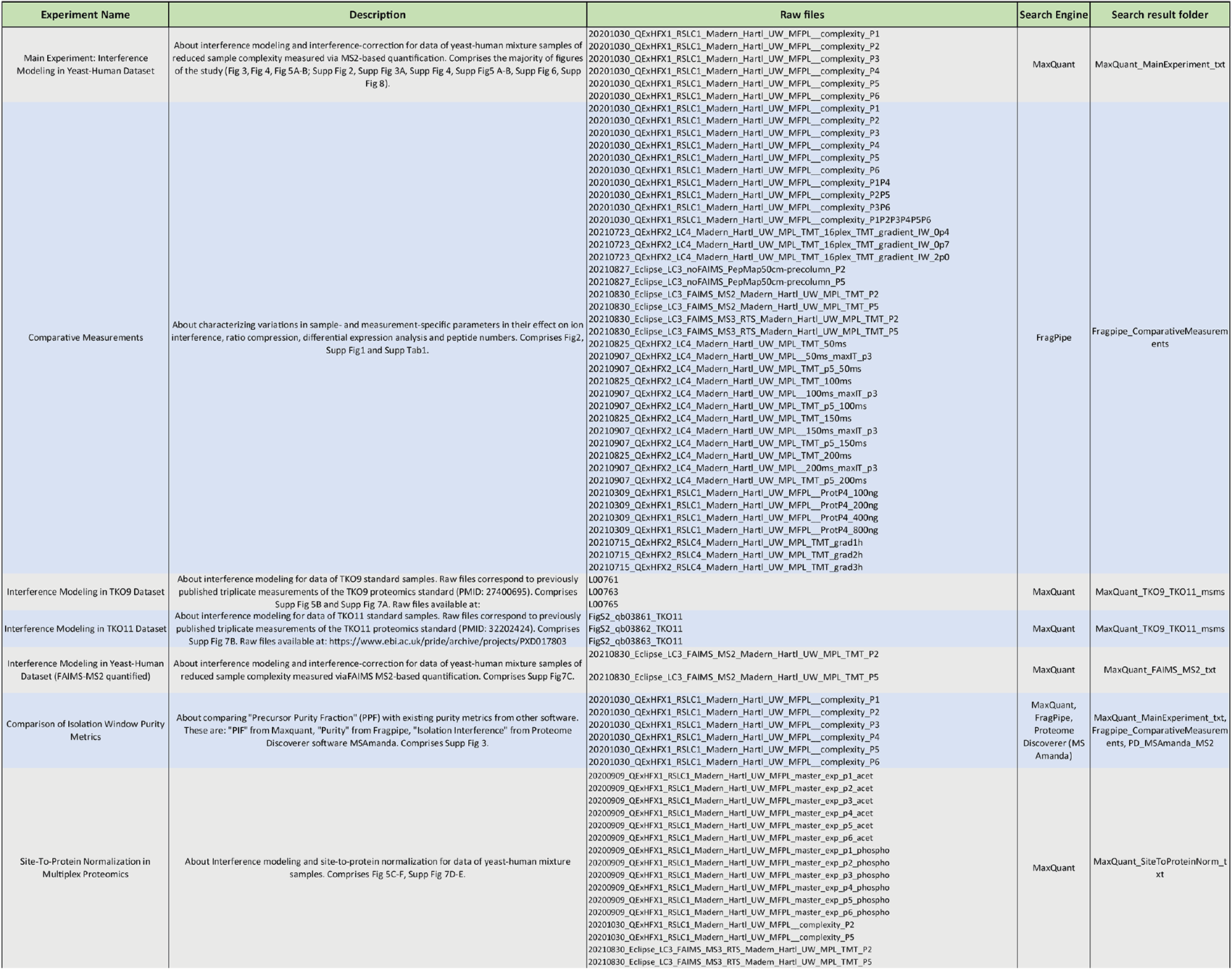
Designation of MS raw files to experiments. Table presents a detailed overview of all measurement raw files and associated database search results (available on PRIDE) and their contribution to the individual results of this study.

